# Field EPSPs of dentate gyrus granule cells studied by selective optogenetic activation of hilar mossy cells in hippocampal slices

**DOI:** 10.1101/2024.11.08.622679

**Authors:** Hannah L. Bernstein, Yi-Ling Lu, Justin J. Botterill, Áine M. Duffy, John J. LaFrancois, Helen E. Scharfman

## Abstract

Glutamatergic dentate gyrus (DG) mossy cells (MCs) innervate the primary DG cell type, granule cells (GCs). Numerous MC synapses are on GC proximal dendrites in the inner molecular layer (IML). However, field recordings of the GC excitatory postsynaptic potential (fEPSPs) have not been used to study this pathway selectively. Here we describe methods to selectively activate MC axons in the IML using mice with Cre recombinase expressed in MCs. Slices were made after injecting adeno-associated virus (AAV) encoding channelrhodopsin (ChR2) in the DG. In these slices, we show that fEPSPs could be recorded reliably in the IML in response to optogenetic stimulation of MC axons. Furthermore, fEPSPs were widespread across the septotemporal axis. However, fEPSPs were relatively weak because they were small in amplitude and did not elicit a significant population spike in GCs. They also showed little paired pulse facilitation. We confirmed the extracellular findings with patch clamp recordings of GCs despite different recording chambers and other differences in methods. Together the results provide a simple method for studying MC activation of GCs and add to the evidence that this input is normally weak but widespread across the GC population.

**KEY POINTS:**

- We describe a method to activate the MC input to GCs selectively using optogenetics in hippocampal slices
- MC excitation is weakly excitatory but so common among GCs that a field EPSP is generated at the site of MC synapses on GCs
- MC excitation of GCs is consistent across the septotemporal axis and contralaterally
- Using the characteristics of optogenetically-evoked fEPSPs, electrical stimulation of the MC input to GCs can be optimized.

## INTRODUCTION

The DG has been suggested to play an important role in many cognitive and behavioral functions and contributes to diverse pathological conditions (Buckmaster and Schwartzkroin, 1994; Sloviter, 1994; Ratzliff et al., 2002; Scharfman, 2016). Despite its important roles, there are many questions about DG, and one is the role of mossy cells (MCs). MCs are a major cell type of the hilus of the DG, are glutamatergic, and project primarily to the proximal third of the molecular layer (the inner molecular layer; IML) where they form a dense projection to GCs (Amaral et al., 2007; Scharfman and Myers, 2012).The IML axon terminals are primarily in parts of the DG that are far from the MC cell bodies of origin, and mostly contact GC dendrites in the IML (Ribak et al., 1985; Buckmaster et al., 1996; Scharfman and Myers, 2012). However, MCs also innervate inhibitory neurons in the GC layer that have dendrites in the IML (Sloviter, 1994; Buckmaster et al., 1996). MCs also have local axon collaterals in the hilus, where they excite GABAergic neurons (Scharfman, 1995; Buckmaster et al., 1996; Larimer and Strowbridge, 2008). The inhibitory neurons activated by MCs innervate GCs, leading to the ability of MCs to inhibit GCs indirectly.

The purpose of this study was to develop a method to study the MC input to GCs selectively using extracellular recordings in slices. Extracellular recording is commonly used to study other highly organized, lamellar inputs to hippocampal principal cells, and provides a way to examine subsets of synchronized principal cells rather than individual neurons. While single cell recordings are useful, understanding how synchronized subpopulations behave is also important. This is highly relevant to MC excitation of GCs because MCs form en passant synapses in the IML, potentially synchronizing subsets of GCs (Buckmaster et al., 1992; Buckmaster and Schwartzkroin, 1994; Buckmaster et al., 1996; Wenzel et al., 1997). Furthermore, extracellular recordings are easier than intracellular recordings and therefore a faster way to obtain information about the MC input to GCs for a non-specialist.

Understanding MC effects throughout the septotemporal axis is an important issue because MCs project throughout the septotemporal axis and contralaterally (Amaral et al., 2007; Scharfman and Myers, 2012). Moreover, dorsal MCs and ventral MCs differ in their projection to the distal DG. Ventral MCs have an axon that is restricted to the IML in the dorsal DG, but dorsal MCs axons terminate in all parts of the ML of ventral DG (Botterill et al., 2021; Houser et al., 2021). It was shown that stimulation of the ventral axons of dorsal MCs synergized with the perforant path input (Houser et al., 2017), potentially explaining a strong role of ventral MCs in behavior (Fredes et al. 2020, 2021). Also, ventral MCs in slices showed more burst firing than dorsal MCs when exposed to pharmacological agents that increased excitability (Jinno et al. 2003). Ventral MCs have greater expression of calretinin (Blasco-Ibanez and Freund, 1997; Fujise et al., 1998). Other differences in dorsal and ventral MC expression also have been reported (Cembrowski et al., 2016).

To selectively activate the MC input to GCs we used optogenetics. Two mouse lines with Cre recombinase (Cre) in MCs were used and a virus encoding channelrhodopsin (ChR2) and enhanced yellow fluorescent protein (eYFP) was injected into one location, either the dorsal or ventral DG. Slices were made throughout the septotemporal axis bilaterally in the same animal. We found fEPSPs could be consistently recorded in response to an optogenetic stimulus of the IML. The fEPSPs were greatest in the IML and appeared to be generated there. Interestingly, all fEPSPs were small and population spikes were difficult to evoke. The weak fEPSPs were present at all levels of the septotemporal axis and both blades of the DG, suggesting they were robust despite their weak strength. The results were confirmed by whole-cell recording from GCs. Together the data provide a useful and relatively simple approach to study MC effects on GCs and support the consensus from previous studies that the MC input to GCs is typically weak. The results suggest, however, that the weak effect is remarkably consistent throughout the MC projection.

## METHODS

All procedures were reviewed and approved by the Institutional Animal Care and Use Committee of The Nathan Kline Institute. Experiments were performed in accordance with guidelines of the Nathan Kline Institute and national guidelines for the care and use of laboratory animals. All chemicals were purchased from Sigma-Aldrich unless otherwise specified.

### I. Terminology

For the purposes of this study, dorsal or septal DG refers to the part of the DG that is near the dorsal aspect of the forebrain and is closest to the septum (Fig. 1A-B). Ventral or temporal DG refers to the part of the DG near the temporal pole of the septotemporal axis (Fig. 1A-B). The term upper blade refers to the blade that is closest to area CA1 (Fig. 1C), also called superior or suprapyramidal blade. The lower blade refers to the blade furthest from area CA1 (Fig. 1C), also called the inferior or infrapyramidal blade. The crest or apex refers to the point where the blades meet (Fig. 1C).

**Figure 1.**
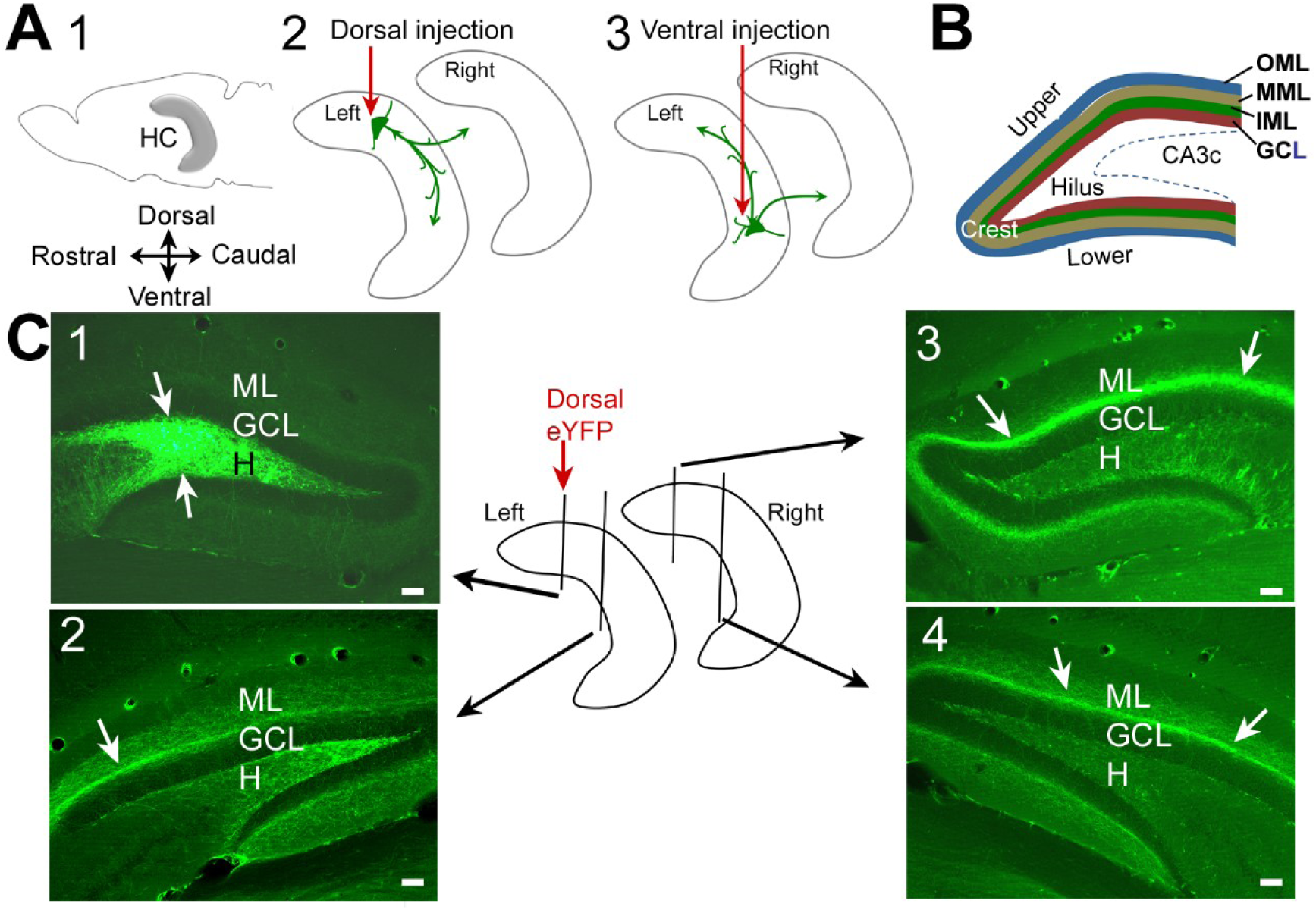
Experimental approach and expression of AAV2-eYFP in *Drd2-*Cre mice. **A** 1. A schematic of the mouse brain illustrates the location of the hippocampus (HC) in gray. Dorsal is up; Rostral is to the left. 2. A diagram illustrates the injection of AAV either in dorsal (2) or ventral (3) hippocampus. The axon of a dorsal MC (green arrows) reflects the major projection, which is to the ipsi- and contralateral IML. **B.** The DG layers are shown: outer molecular layer (OML); middle molecular layer (MML); inner molecular layer (IML); granule cell layer (GCL); hilus. CA3c is the area CA3c pyramidal cell layer. The term “ML” in part C and elsewhere refers to all three ML subdivisions (OML, MML, IML). **C.** In the center is a schematic of the injection of AAV2-eYFP into the dorsal DG and where sections were selected for 1-4. 1. Expression at the injection site was in the hilus. 2. Expression distal to the injection site in the same hemisphere show staining in the IML (arrows). 3. Expression contralateral to the injection site show staining in the IML (arrows). 4. Expression in the IML contralateral to the injection site, at an anterior-posterior level similar to 3 (arrows). Calibrations, 50 µm.

### II. Animals and genotyping

One mouse line expressed Cre in cells that have the dopamine receptor type 2 (Drd2), which, in the DG, are primarily localized to MCs (Gangarossa et al., 2012; Puighermanal et al., 2015). It has been reported that some GABAergic neurons express Cre in the mice (Puighermanal et al., 2015; Azevedo et al., 2019), so another mouse line with Cre in cells that have the calcitonin receptor-like receptor (Crlr; (Jinde et al., 2012)) was also used. The *Drd2*-Cre mice and *Crlr*-Cre mice were graciously provided by Drs. E. Valjent and Dr. K. Nakamura respectively. Hemizygous *Drd2*-Cre males were bred to C57BL/6NCrl females (#027, Charles River Laboratories). Hemizygous *Crlr-*Cre males were bred in-house to C57BL/6J female mice (#000664, Jackson Laboratories). Hemizygous *Drd2*-Cre or *Crlr*-Cre offspring were used for experiments.

Breeders were provided a nestlet (2”x2”; W.F. Fisher) and fed Purina 5008 chow (W.F. Fisher). After weaning at 25-30 days of age, mice were housed with others of the same sex (2-4/cage) and fed Purina 5001 chow (W.F. Fisher). At all times, mice were provided food and water *ad libitum*. They were housed in standard mouse cages with a 12 hr light:dark cycle (lights on: 7:00 a.m.-7:00 p.m.) and relative humidity between 64-78%. Mice were housed overnight in the laboratory (2-4/cage) to acclimate them prior to slice preparation the next morning. Tail samples were frozen and genotyped later (Mouse Genotyping Core, New York University Medical Center).

### III. Viral expression of channelrhodopsin (ChR2)

#### A. Viruses

The viruses were adeno-associated virus (AAV)-Ef1a-DIO-hChR2(H134R)-eYFP (serotype 2 or 5) to express humanized ChR2 tagged with eYFP in Cre-expressing cells. Titers for all injected viruses were ∼10^12^ vector genomes (vg) per ml. Aliquots of 5 µl of each virus were stored at −80°C until the day of use. Viruses were obtained from the Karl Deisseroth laboratory or the University of North Carolina Vector Core or Addgene.

#### B. Surgical injection of viruses

Animals (6-8 weeks old) were anesthetized with 4% inhaled isoflurane in an induction chamber (#V711801; SurgiVet), then transferred to a rodent stereotaxic apparatus (#502063, World Precision Instruments; WPI). Body temperature was maintained at 37°C with a homeothermic heating pad (#507220F, Harvard Apparatus) for the duration of surgery. Isoflurane (1-2%; Aerrane, Piramal Enterprises) mixed with oxygen was delivered through a nose cone (WPI). Buprenex (buprenorphine; 0.1 mg/kg, subcutaneous; Reckitt-Benkiser Pharmaceuticals) was administered shortly after the onset of anesthesia to reduce discomfort. The skin of the head was shaved, sterilized with Betadine (Purdue Products) and 70% ethanol (Pharmaco-Aaper), and then an incision was made. A hand drill (#177001, Dremel Instruments) was used to create a craniotomy above the injection site. Using a nanoliter-precision automated syringe pump (#UMP3, WPI) with a syringe (#Nanofil, WPI) and 33-gauge needle (#Nanofil-10F, WPI), 150 nl of virus was injected into hippocampus at 100 nl/min. The coordinates (in mm from Bregma) for dorsal injections were: Anterior-Posterior (A-P) 2.0, Medio-Lateral (M-L) 1.3, depth 2.1 and for ventral injections they were A-P 3.4, M-L 2.6, depth 2.4. For experiments using whole cell recording, there were some mice that were injected both in dorsal and ventral DG to maximize viral expression, and slices were made contralaterally in these cases. The coordinate for ventral expression was a depth of 2.8 instead of 2.4 because that appeared to increase the ability to target ventral MCs. The needle was left at the injection depth for 5 minutes to allow diffusion of the virus before being slowly retracted. The incision was closed with Vetbond (3M Inc.). Animals were transferred to a clean cage and observed until they recovered from anesthesia. They were housed alone until use.

### IV. Evaluation of expression

Animals were deeply anesthetized with isoflurane (by inhalation in a closed glass jar) followed by urethane (2.5 g/kg, intraperitoneal or i.p.). The abdominal cavity was opened and the animal was transcardially perfused with ∼10 ml 0.9% NaCl in distilled (d)H_2_O followed by 20 ml 4% paraformaldehyde in 0.1 M phosphate buffer (PB; pH 7.4), using a 25 gauge butterfly needle attached to a peristaltic pump (#Minipuls 2, Rainin). Brains were removed and postfixed overnight at 4°C and then cut into 50 µm coronal sections using a vibratome (#TPI1000, Ted Pella, or VT1000P, Leica). Sections were collected into cryoprotectant (25% glycerol, 30% ethylene glycol, 45% 0.1M phosphate buffer) and stored at 4°C.

Immunofluorescence was conducted as previously described (Duffy et al., 2013). Free-floating sections were washed in 0.1 M PB (3 times, 5 min each) and transferred to blocking serum (10% normal donkey serum for parvalbumin (PV); 10% normal goat serum for GluR2/3 and neuropeptide Y (NPY), and 0.25% Triton-X 100, diluted in 0.1 M PB) for 1 hour. Sections were then transferred directly to primary antibody solution (0.25% Triton X-100 diluted in 0.1 M PB), incubated with primary antibody (mouse monoclonal to PV (1:500; #MAB1572, Millipore), rabbit polyclonal to GluR2/3 (1:100; #AB1506, Millipore) or NPY (1:1000; #T-4070, Peninsula), and gently agitated on a rotator overnight at 4°C. The next day, sections were washed in PB (3 times, 5 min each) and then incubated with secondary antibody (AlexaFluor 546 Donkey anti-mouse for PV, goat anti-rabbit for GluR2/3 or NPY, both 1:400, Thermo Fisher Scientific Inc.) in secondary antibody solution (1% normal donkey serum for PV or 1% normal goat serum for GluR2/3 or NPY, and 0.25% Triton-X 100, diluted in 0.1 M PB). After washing in PB (3 times, 5 min each), sections were mounted and allowed to dry at room temperature overnight. Slides were coverslipped using ProLong Gold antifade solution (Thermo Fisher) and stored at 4°C.

Stained sections were imaged on a laser scanning confocal microscope (#LSM 510; Zeiss Inc.). Double fluorescent labeling was quantified using LSM ImageBrowser software (Zeiss). Double labeling was determined by acquiring z-stack images at 1.5-2.0 µm intervals of a cell and locating the brightest optical section of the cell for each color channel. A cell was considered double labeled if the brightest optical sections of a cell were at the same depth. Cells that appeared to be double labeled but were actually overlying one another could be discriminated with this method and an example is shown in Supplemental Figure 1D.

### V. Electrophysiology

#### A. Extracellular recording

Mice were rapidly anesthetized with isoflurane by inhalation. While deeply anesthetized, mice were decapitated, the brain rapidly dissected and placed in a 4-8°C artificial cerebrospinal fluid (aCSF) with sucrose instead of NaCl (in mM: 252 sucrose, 3.5 KCl, 2.0 MgSO_4_-7H_2_O, 2.4 CaCl_2_, 1.25 NaH_2_PO_4_, 26 NaHCO_3_, 10 D-glucose, pH 7.4). After 1-3 min, the brain was blocked to cut it transversely or horizontally. For transverse sections, slices were cut at an approximately 15° angle to the coronal plane so most of the dorsal hippocampus would be cut along the transverse or lamellar axis. The extreme septal pole was excluded from use because the upper DG blade is abbreviated. For horizontal sections, the dorsal surface of the brain was trimmed so it would lie flat and horizontal sections were cut from the ventral hippocampus until the sections were outside the transverse axis. The extreme pole was not used because it is outside the transverse axis. We acknowledge that the use of the term transverse is approximate.

For all sectioning, the brain was glued to a metal plate using cyanoacrylate. The metal plate was placed in a holding container filled with chilled sucrose-based aCSF and cut (400 µm thick) using a vibratome (#Microm HM 650V, Richard-Alan Instruments). Sections containing hippocampus and adjacent regions were trimmed and then placed in room temperature sucrose-based aCSF in a beaker with carbogen (5% CO_2_, 95% O_2_) for 1-5 min. Slices were transferred to an interface-style recording chamber (made in-house; Scharfman 2001). Using this chamber, all slices could be maintained under similar conditions immediately after the dissection until the end of the experiment when they were removed and fixed by immersion in 4% paraformaldehyde in 0.1M Tris buffer.

Slices were placed on a nylon mesh, aCSF entered from below, and aCSF flowed up and over the mesh. Slices were placed on a nylon mesh, aCSF entered from below, and aCSF flowed up and over the mesh. A thin layer of aCSF was over the mesh. The height of the fluid was adjusted by a piece of a Kimwipe that was draped over the edge of the mesh. It led to a distant port where there was gentle suction. The degree that the Kimwipe was on the mesh led to more or less fluid over the mesh. A greater area of the Kimwipe on the mesh lowered the height of the fluid. When the Kimwipe was barely on the mesh at all, the height of the fluid on the mesh was greatest. This arrangement kept slices submerged up to their upper surfaces. This arrangement kept slices submerged up to their upper surfaces. Warmed (32°C) humidified air was vented over the surfaces. Slices were in the chamber for 30 min with sucrose-based aCSF at 1-2 ml/min (#Minpuls 2, Gilson) and then NaCl-based aCSF (125 mM NaCl instead of sucrose) was used. Recordings began at least 30 min after the switch from sucrose-based aCSF to NaCl-based aCSF and terminated up to 6 hrs later.

The interface chamber does not allow visualization of DG neurons to judge their health so slices were initially screened for viability using field potential recordings of the response to an electrical stimulus to the PP. For PP stimulation, a stimulating electrode was placed on the slice surface at the point along the hippocampal fissure just below the subiculum where the PP is a white fiber bundle as it enters the DG. Field recordings were made in the GC layer (GCL) at the border with the hilus, either near the crest, or near the center of the upper blade, approximately 250 µm from the stimulating electrode. To assess viability, the population spike was not used because the GCs do not generate many APs normally, so the population spike is normally small. The amplitude of the positivity, upon which the population spike is superimposed, was used instead, since past studies found that when the positivity is >3 mV and the population spike is small (<2 mV) many GCs can be recorded intracellularly with characteristics consistent with healthy cells (Scharfman, 1991, 1994; Scharfman et al., 2000; Scharfman et al., 2002).

Monopolar electrodes used for stimulation were made from 35 µm-diameter Teflon-coated stainless-steel wire (A-M Systems Inc.) and triggered in pClamp (Molecular Devices) using 50 µA intensity, 10-100 µsec duration square pulses, generated from a stimulus isolation unit (#Isoflex, AMPI Instruments).

Electrodes used for extracellular recording were 1-5 MΩ and filled with NaCl-based aCSF. Electrodes were pulled from 1.2 mm o.d., 0.85 mm i.d. borosilicate glass with an internal filament (WPI) using a horizontal puller (#P-87, Sutter Instruments). Signals were digitized at 10 kHz for field and 30 kHz for intracellular recordings (#Axoclamp2B and Digidata 1440A, Molecular Devices).

The latency to onset was measured from the end of the light pulse to the fEPSP onset. The fEPSP slope was the maximum slope of the rising phase. The amplitude was measured from the pre-stimulus baseline to the peak.

#### B. Whole-cell recording

Whole-cell recording was used to confirm what had been found using fEPSPs and intracellular recordings. Whole-cell recordings used a different recording chamber and arrangement for optogenetics, and other aspects of the methods differed from extracellular recordings. Although this was a limitation of the comparison, it also had advantages, in that it allowed us to determine if there would be confirmation despite differences in methods. If so, the data would make a better case that the results from field potentials could generalize to other recording methods.

Slices were prepared with methods that are typical for whole cell recordings. The sucrose aCSF contained (in mM): 90 sucrose, 80 NaCl, 2.5 KCl, 1.25 NaH_2_PO_4_, 25 NaHCO_3_, 10 D-glucose, 4.5 MgSO_4_-7H_2_0, and 0.5 CaCl_2_. Slices were 350 μm thick instead of 400 μm. After slice preparation, slices were placed in a holding chamber (made in-house) in oxygenated (95% O_2_, 5% CO_2_) sucrose-based aCSF. The holding chamber consisted of a beaker containing an insert with a nylon mesh where slices were placed without overlap. An aerator was placed 1″ below the netting so fine bubbles of 95%O_2_, 5%CO_2_ could be directed towards the slices. The beaker was filled with aCSF so that the slices were submerged fully. The beaker was placed in a water bath (Poly Pro Water Bath; Revolutionary Science), and the temperature was increased gradually to 35°C over the first 15-20 min and maintained at 35°C thereafter. After slices had been in the holding chamber for 45 min, slices were kept in sucrose-based aCSF at room temperature until use.

Slices were placed in a recording chamber (Model #RC-27LD; Warner Instruments). ACSF entered from the side and flowed above and below the slice so it was entirely submerged. Slices were held down with a nylon mesh (Item #SS-4-500H; Warner). ACSF was removed on the other side of the chamber from the entry pport bu gentle suction. Slices were warmed to 32°C with an in-line heater (Item #SH-27B; Warner), and perfused at 6-7 mL/min by a peristaltic pump (#Masterflex C/L; Cole-Parmer) on an infrared differential interference microscope (#BX51WI; Olympus). Slices were perfused with NaCl-aCSF containing the same constituents as described above except 1) 130 mM NaCl was used instead of 126 mM for extracellular recordings, and 2) 2.5 mM KCl was used instead of 3.5 mM.

Recording electrodes were pulled from borosilicate glass without a filament (1.5 mm outer diameter and 0.86 mm inner diameter, Sutter Instruments) using a micropipette puller (#P-97; Sutter) with tip resistance between 5-10 MΩ. Internal solution was filtered (#0.02 µm, Whatman Anotop 10 syringe filter; Fisher) and contained (in mM): 130 potassium gluconate, 4 KCl, 2 NaCl, 10 4-(2-hydroxyethyl)-1-piperazine ethanesulfonic acid (HEPES), 0.2 ethylene glycol-bis(β-aminoethyl ether)-N,N,N′,N′-tetraacetic acid (EGTA), 4 adenosine triphosphate magnesium salt, 0.3 guanosine triphosphate tris salt, 14 di-tris phosphocreatine, 0.2-0.5% biocytin, and was adjusted to pH 7.25 using 1M potassium hydroxide. Final osmolarity was 280-290 mOsm without biocytin. Aliquots were frozen (−20°C) and used within 6 months.

Recordings were analyzed offline using Clampfit (v. 10.7 or 11.1; Molecular Devices). Resting membrane potential (RMP) was defined as the difference between the MP during recording (without current injection) and the extracellular potential after the recording. Latency to onset and amplitude were measured as described above. Rise time was the slope measured during the time when the EPSP was 10-90% of its peak amplitude. Duration was measured as half-duration, the time from the EPSP onset to the point on the decay when the EPSP reached half its peak amplitude.

### VI. Optogenetics

A 200 µm-diameter fiber optic cable (#M86L01l, Thorlabs) was modified by removing the ferrule and rubber coating from the tip, stripping approximately 2 cm of covering (cladding), and polishing the bare fiber tip with fine emery cloth. The fiber tip was approximately 150 µm thick and threaded through a glass Pasteur pipette with the tip extruding approximately 2 cm. The Pasteur pipette was held by a micromanipulator (#NMN-21, Narashige) so that it was approximately 3 cm from the slice surface (for extracellular and sharp recordings). The glass holder was slanted 15-20° from vertical. For whole-cell recording, the light fiber was positioned 55° from vertical (or 35° from horizontal). Blue light was generated by attaching the fiber to a 473 nm DPSS laser (#LRS-0473, Laserglow), after power was adjusted to 10 mW with a power meter (#PM100D, Thorlabs). Light pulses (typically <2 msec duration) were triggered 1/min or in pairs 10-200 msec apart (1 pair, every 1-2 min).

For both the interface and submerged slice chambers, optogenetics methods were almost identical. The difference was that the fiberoptic tip was in the air above the slice for the interface chamber and had a submerged tip for whole-cell recording. For the interface chamber, the fiberoptic tip was held at an angle by a micromanipulator and the tip was 1 mm above the slice surface, pointing to a location approximately 200 µm from the recording electrode. For the latter, the fiberoptic was held at an angle with a micromanipulator and placed on the IML approximately 200 µm of the recording electrode.

### VII. Pharmacology

The AMPA receptor antagonist 6,7-Dinitroquinoxaline-2,3(1H,4H)-dione (DNQX disodium salt; Tocris) or 6-Cyano-7-nitroquinoxaline-2,3-dione (CNQX disodium salt, Sigma) or GABA_A_ receptor antagonist bicuculline methiodide (Sigma) were dissolved in 0.9% NaCl to make a 10 mM stock solution, stored at −20°C and diluted in aCSF to reach the final concentration before use.

### VIII. Statistics

Results are expressed as mean ± standard error of the mean (SEM). Data are available at Open Science Framework (www.osf.io; Project name, Bernstein et al.; DOI 10.17605/OSF.IO/Z84GS). Power analysis was used to determine if sample size was adequate, for α = 0.05 and 80% power (Statmate2; GraphPad). Statistics were conducted using Prism (GraphPad). Comparisons of two groups were made using Student’s t-test (unpaired, two-tailed) and one- or two-way Analysis of variance (ANOVA) or repeated measures ANOVA (RMANOVA) for more than two groups. For ANOVAs, Tukey-Kramer *post-hoc* tests were conducted to assess statistical differences between groups with correction for multiple comparisons. Interactions and main effects are not reported for ANOVAs if they did not reach significance. Nonparametric tests were conducted when data were not normally distributed by a Shapiro-Wilk’s test or there was heteroscedasticity of variance by Bartlett’s test that could not be corrected by transformation of the data. Non-parametric tests of two groups used a Mann-Whitney *U* test and for >2 groups, a Kruskal-Wallis test was conducted.

## RESULTS

### I. Specificity of Drd2-Cre mice and Crlr-Cre

#### A. *Drd2*-Cre mice

It was important to verify specificity of the *Drd2*-Cre hemizygous mice (referred to as *Drd2*-Cre, below) because a previous description of these mice indicated there was Cre expression in a subset of hippocampal interneurons (Puighermanal et al., 2014) which was also shown later (Azevedo et al., 2019). However, in the latter study, the volume of virus was 500 nl, more than we used. To evaluate expression following viral injection, 5 *Drd2*-Cre mice were injected with AAV2-EF1a-DIO-eYFP (Fig. 1A-B) and euthanized 10-18 days later to examine eYFP expression. As shown in Fig. 1C1, the hilus was labeled extensively at the injection site in the DG. Sections in or near the injection site showed strong eYFP-labeling of neuronal somata in the hilus, with few somata outside the DG, consistent with a Cre line specific for MCs. Sections that were distal to the injection site showed prominent labeling of fibers in the IML (Fig. 1C2-4). The weak IML labeling near the injection site and strong IML labeling distal to the injection site is consistent with the MC axon projection in the distal IML primarily (Buckmaster and Amaral, 2001).

To address specificity further, eYFP-expressing cells were double-labeled using an antibody to the 2/3 subunit of the ionotropic glutamate receptor because it labels MCs (GluR2/3; Leranth et al., 1996). Mice were either injected in dorsal (n=3) or ventral DG (n=2). An example from a mouse injected dorsally is shown in Supplemental Fig. 1. For quantification, sections were selected from the injection site and areas next to it (5-7 sections/mouse, 300 µm apart). In the mice injected in dorsal hippocampus, 538/545 (98.6%) of hilar eYFP-expressing cells also expressed GluR2/3 (Supplemental Fig. 1A-B). For the mice injected ventrally, 661/695 (94.8%) cells were double-labelled. For these analyses, the hilus was defined as zone 4 of Amaral (Amaral, 1978).

In contrast to double-labeling of eYFP and GluR2/3, no eYFP-expressing cells were labeled using an antibody to parvalbumin (PV; Supplemental Fig. 1C) which stains a major subclass of DG GABAergic neurons called basket cells or perisomatic targeting cells; (Sloviter, 1994; Freund and Buzsaki, 1996; Houser, 2007; Hosp et al., 2014). It was extremely rare to find eYFP-expressing cells double-labeled using an antibody to NPY, which is a marker of most hilar GABAergic cells, called HIPP cells (***<u>HI</u>***lar cell body and projection to the terminal zone of the ***<u>P</u>***erforant ***<u>P</u>***ath; (Bering et al., 1997; Freund and Gulyas, 1997; Scharfman et al., 2002; Scharfman and Gray, 2006; Sperk et al., 2007); Supplemental Fig. 1D). Double-labeling occurred in 4/337 (1.2%) of hilar cells that also expressed eYFP (for dorsal injections) and 9/328 (2.7%) for ventral injections. These data suggest that there were up to 3% of cells that showed expression and were not MCs in *Drd2*-Cre mice.

#### B. *Crlr-Cre* mice

For Crlr-Cre mice, expression of eYFP was similar to Drd2-Cre mice when virus was injected in dorsal hippocampus (Supplemental Fig. 2A1-3). However, ventral injections led to labeling of a subset of CA3 PCs in CA3b and c (Supplemental Fig. 2B1-3). The labeling of some CA3 PCs was noted in a prior report (Jinde et al., 2012) but it was not clear whether this varied across the septotemporal axis. We only saw the CA3 labeling ventrally (Supplemental Fig. 2B1-3). To determine if there was sufficient expression in CA3 to produce a functional effect, tests were made in slices. As shown in Supplemental Figure 2A4-5 and 2B4-5, in dorsal slices there were no effects of optogenetic stimulation in CA3 on recordings in CA3 (n=3 slices, 2 mice). However, robust field potentials were recorded in ventral slices (Supplemental Fig. 2B4-5; n=3 slices, 2 mice). These data suggest that optogenetic stimulation evoked PC activation in ventral PCs of Crlr-Cre mice, probably by direct excitation. Therefore, subsequent viral injections in Crlr-Cre mice were only made in dorsal DG.

### II. Optogenetic stimulation of MC axons in the IML evoke a small fEPSP in GCs

There were 26 animals injected with virus encoding ChR2 that were used for extracellular and sharp electrode recordings. They were 80.3 ± 7.6 days old when they were used for slice preparation, and the average time between viral injection and slice experiments was 23.9 ± 2.4 days. The majority of the 26 animals were *Drd2*-Cre (n=18) and the others were *Crlr*-Cre (n=8). Of the 26 mice, 19 received dorsal injections and the 7 others had ventral injections. All Crlr-Cre mice had dorsal injections, for reasons mentioned in the preceding section.

Preliminary experiments tested whether any subfield of the hippocampal slice showed a response to light using extracellular recording and *Drd2*-Cre mice (1-6 slices/mouse, 6 mice). Even long pulses (20 msec) failed to exhibit responses in most locations. The one combination of optic fiber position and recording site that was successful used light directed at the IML with a recording position that was in the IML. To avoid direct stimulation, the optogenetic probe was pointed at least 250 µm from the recording site. With these optogenetic stimulus and recording positions, a brief stimulus could elicit what appeared to be a fEPSP in the IML. In the GCL and MML there was a similar but much smaller response (Fig. 2). Therefore, the ability to focus light in a small area seemed clear, despite the fact that the light was likely to scatter outside the small area.

**Figure 2.**
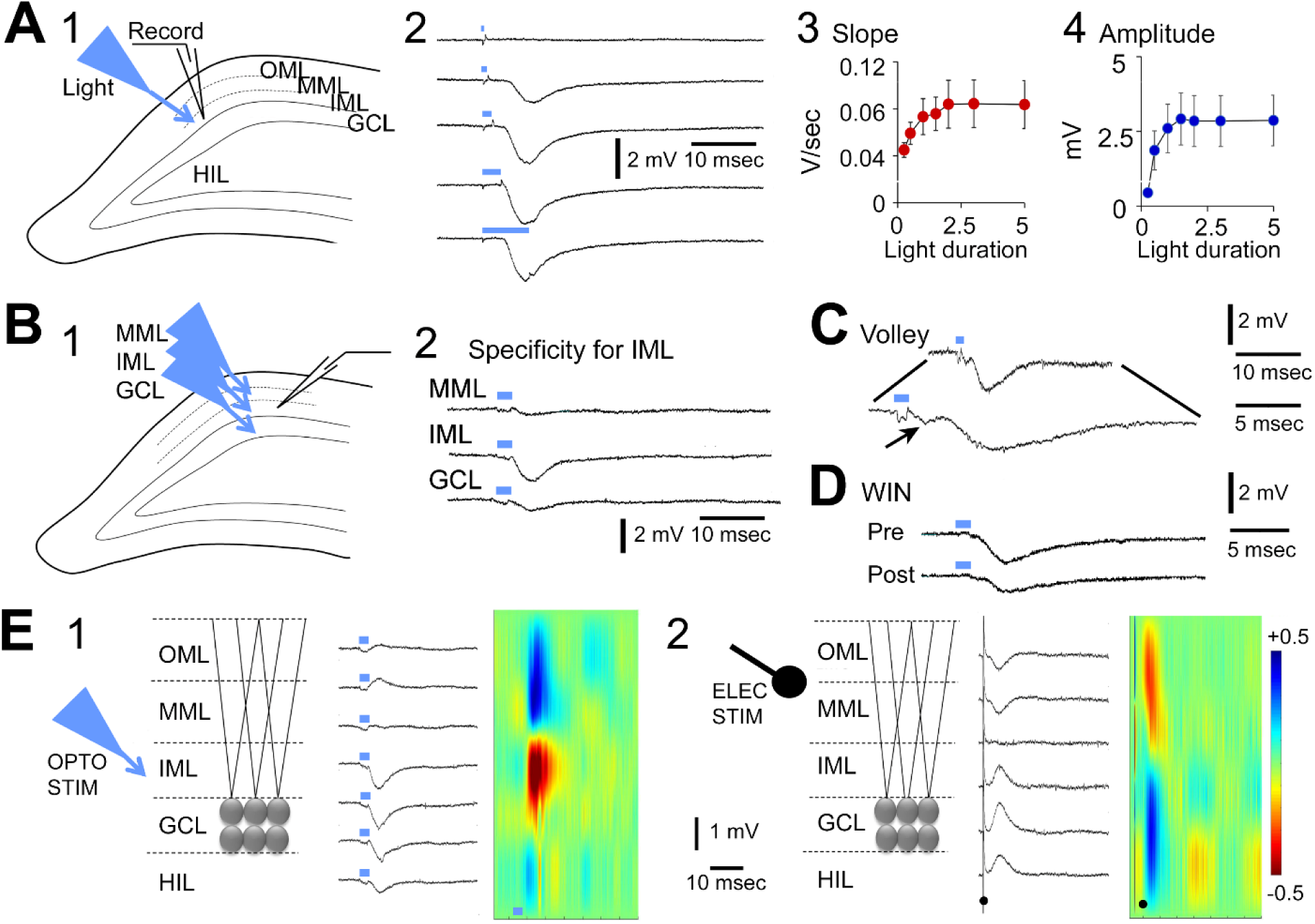
Optogenetic stimulation of the IML evokes field EPSPs (fEPSPs) in the IML. **A.** *Drd2*-Cre or *Crlr*-Cre mice were used after a single injection of AAV2-ChR2 in either dorsal (*Drd2*-Cre, *Crlr*-Cre) or ventral (*Drd2*-Cre) DG. Field potentials evoked by light were recorded. 1. The experimental arrangement is shown. Light was directed to the IML at least 250 µm from a recording electrode in the IML. 2. Examples of responses to increasing durations of light pulses (blue bars). There was a gradual increase in response amplitude, consistent with a fEPSP. Note that after reaching a maximum amplitude, increasing the light pulse duration further did not lead to a greater response. 3. The graded nature of the response to light is shown for fEPSP slope and amplitude in mice tested with the same light pulse durations (n=11 slices, 1 slice/mouse). **B.** FEPSPs are specific to the IML. 1. The experimental preparation is shown. The recording site was in one location in the IML while the light fiber was moved. The light pulse duration was the same for all recordings. 2. The fEPSP was largest when the light fiber pointed to the IML and was greatly reduced when the light fiber was moved so it was pointed at the GCL or MML. **C.** Example of a fEPSP preceded by a fiber volley (arrow) indicating the optogenetic stimulation activated axons in the IML. The volley was not elicited by light directed at the GCL or MML (not shown), suggesting specificity for IML axons. The same trace is shown at two different timescales. **D.** A fEPSP evoked by an optogenetic stimulus to the IML is shown before and after exposure to WIN 551212, an antagonist of MC synapses on GCs (Chiu and Castillo, 2008). **E.** Current source density (CSD) analysis. The recordings in 1 and 2 are from the same slice. 1. CSD analysis of the optogenetically-evoked fEPSP. *Left:* A diagram shows the layers of the DG where recordings were made in response to light activation of the IML. *Center:* Responses evoked a fEPSP that was largest in the IML. *Right:* Current source density (CSD) analysis showed a current sink (red) in the IML, consistent with preferential activation of MC axons. 2. CSD analysis of responses to electrical stimulation of perforant path (PP) axons.. *Left:* A diagram shows the site of the stimulating electrode, with the tip in the location where PP fibers terminate. *Center:* Stimulation evoked fEPSPs that were maximal in the OML. *Right:* The CSD showed a current sink in the OML, consistent with a PP-evoked fEPSP.

The characteristics of the response supported the idea that it reflected EPSPs in GCs. Thus, responses were graded like an EPSP, i.e. increasing the duration of the light pulse led to a graded increase in slope and amplitude (Fig. 2A1-4). The fEPSP had a rapid onset, usually beginning at the end of the light pulse or shortly thereafter (Fig. 2A2). To determine the mean, responses were chosen that were between 50 and 100% of the maximum response. The mean latency, measured from the end of the light pulse, was 0.69 ± 0.11 msec (n=42 slices, 16 mice). The mean latency, from the start of the pulse, was less than 3 msec because the light pulses were less than 2 msec long. These latencies suggest that the fEPSP reflects a monosynaptic pathway (MC → GC). Characteristics of latency, slope and amplitude of fEPSPs are shown in Supplemental Table 1.

Consistent with the idea that fEPSPs were excitatory, the AMPA receptor antagonist CNQX) 10 µM) decreased the fEPSP (to 24.9 ± 4.8% of control after 30 min, n=3 slices, 1 slice/mouse; paired t-test, t=5.8, df=2, p=0.028), but the GABA_A_ receptor antagonist bicuculline methiodide did not (BMI; 10 µM; 101.0 ± 1.9% of control at 30 min, n=3 slices, 1 slice/mouse; paired t-test, t=0.78, df=3, p=0.495; Supplemental Fig. 3). The use of bicuculline also allowed us to show that the optogenetic stimulus was able to activate the MC input to GABAergic neurons, which inhibits GCs. This was done by stimulating optogenetically prior to an electrical stimulus to the PP (Supplemental Fig. 3). The optogenetic stimulus decreased the population spike evoked by the PP stimulus (paired t-test, perforant path response with vs without a prior optogenetic stimulus, t=9.00, df=2, p=0.012; n=3 slices, 3 mice; Supplemental Fig. 3). In the presence of bicuculline, there was no difference (paired t-test, t=0.23, df=2, p=0.840; n=the same 3 slices; Supplemental Fig. 3).

Other characteristics of fEPSPs were identified by changing the position of the optic fiber (Fig. 2B1-2). In 18 slices from 12 mice, minor movements of the optic fiber were made either towards the MML or in the other direction, towards the GCL. In each case the fEPSP decreased greatly even by minor movement of the optic fiber (Fig. 2B2). In contrast, movements of the optic fiber within the IML, either further (up to 300 µm) or closer to the recording pipette, had little effect on the fEPSP amplitude (<5% change in amplitude, n=12 slices in 6 mice). The similarity of responses when the light fiber was moved along the IML instead of away from the IML is consistent with the en passant synapses of MCs in the IML (Buckmaster et al., 1992; Buckmaster and Schwartzkroin, 1994; Buckmaster et al., 1996; Wenzel et al., 1997). Specificity of fEPSPs to the IML was also evident from recordings of a fiber volley before the EPSP with recording sites in the IML (10/34 fEPSPs; Fig. 2C) but not the GCL or MML.

To confirm the responses to light were due to activation of MC axons, fEPSPs were tested for sensitivity to WIN 55,212 mesylate (WIN), a cannabinoid receptor type 1 (CB1) agonist which is highly expressed on MC terminals in the IML (Chiu and Castillo, 2008), Exposure to 5 µM (n=2 slices in 2 different mice) or 10 µM (n=1 slice in a 3rd mouse) WIN for 30 min reduced the slope and amplitude of the light-evoked fEPSP to 44.0 ± 4.9% of control in 30 min (Fig. 2D). The fact that the fEPSP was not completely blocked may be due to the weak affinity of WIN for CB1 receptors on glutamatergic terminals relative to non-glutamatergic terminals (Hajos and Freund, 2002).

Specificity of optogenetically-triggered fEPSPs to the IML was also supported by current source density analysis (Fig. 2E1; n=5 slices, 5 mice). Thus, when light was focused on the IML and the recording electrode was moved from the hilus to the hippocampal fissure (along a path oriented perpendicular to the GCL), the fEPSP increased as the IML was approached and reached the maximum in the IML (Fig. 2E1). An ocular micrometer was used to make recording electrode locations a similar distance apart. Using this approach, current source density analysis showed a current sink in the IML (Fig. 2E1; n=3 slices, 1/mouse). In contrast, the response to PP stimulation produced a current sink in the area of the OML or MML, where its synapses are located (Fig. 2E2; n=4 slices, 3 mice).

In 28 slices (7 animals) there was no detectable response to light in the IML or nearby (GCL, MML). The expression of virus was only checked after the recordings were made and the slice was removed and brought to a microscope. In each of these animals the slices had no detectable fluorescence. In addition to these experiments, three mice were used that lacked Cre recombinase and did not have a viral injection. No response to light could be detected (n=5 slices). Even long pulse durations (up to 50 msec) with increased power (up to 50 mW) were ineffective.

We also examined the response recorded in stratum lucidum (SL) of CA3 to the same optogenetic stimulus that was used to elicit the fEPSP in the IML and no fEPSP was recorded in SL (Fig. 3A). These data are consistent with the idea that the EPSP elicited by MCs in GCs is indeed a weak EPSP.

**Figure 3.**
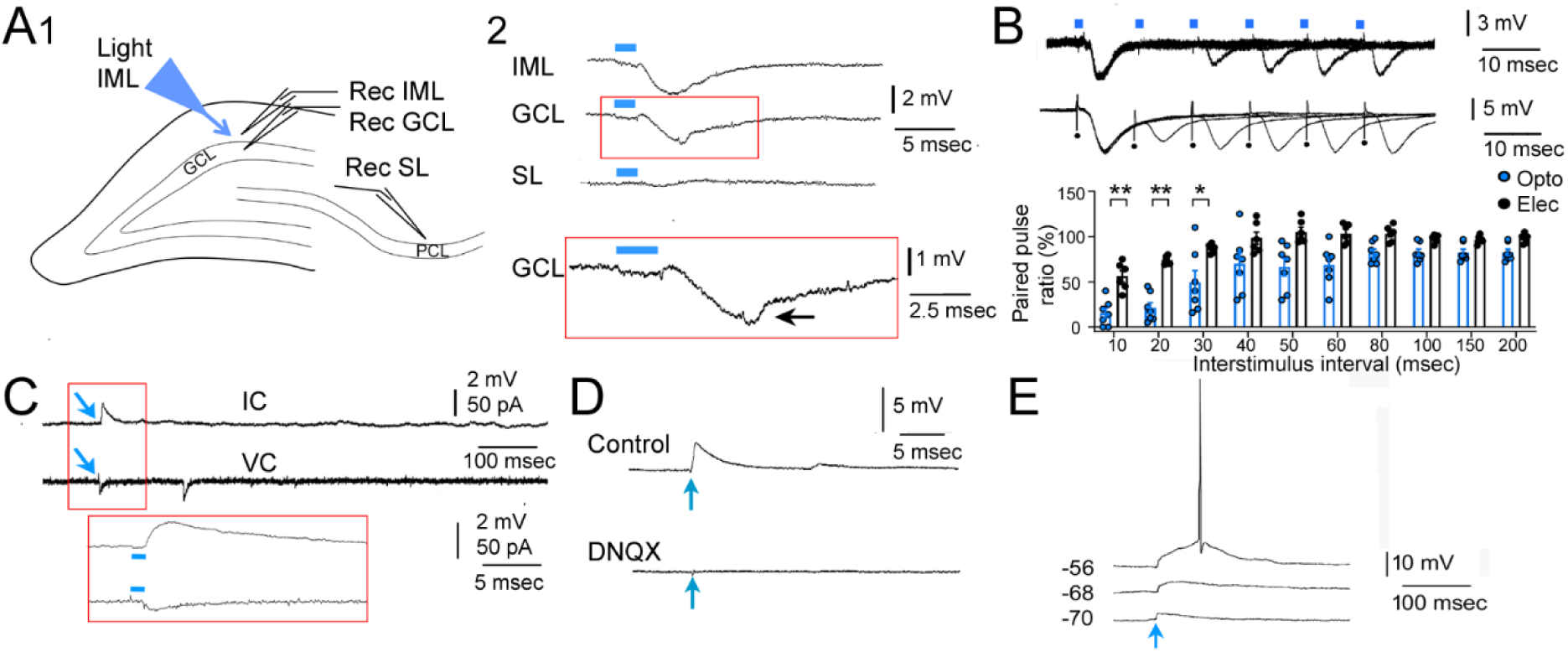
fEPSPs elicited by optogenetic stimulation of IML axons failed to activate CA3. **A.** 1. The experimental arrangement is illustrated. Stratum lucidum, SL. 2. Recordings were made in the IML, GCL, and SL in response to an optogenetic stimulus (blue bar). There was a fEPSP in the IML. In the GCL, the fEPSP was unable to evoke a large population spike. In SL, there was no effect of the optogenetic stimulus, suggesting inability to activate CA3. Inset: The recording in the GCL is expanded to show an extremely small population spike (arrow), consistent with weak excitation of GCs. **B.** *Top*: Pairs of optogenetic stimuli (blue bars) were elicited with 10-200 msec interstimulus intervals and are superimposed. Data are from 6 slices of 7 mice. *Center*: Pairs of electrical stimuli (blue bars) were elicited. Data are from 7 slices of 7 mice. *Bottom*: Comparison of paired pulse ratios show significant differences for 10 (*, p=0.002), 20 msec (p=0.001), and 30 msec (p=0.038) interstimulus intervals (Mann-Whitney *U* tests) *p<0.05; **p<0.01. **C.** Optogenetic stimulation of the IML (blue arrow) in a patched GC led to an EPSP in current clamp (IC). The same cell was then recorded in voltage clamp (VC) and the optogenetic stimulus evoked an inward current. There is also a second current after the evoked inward current which is likely to be a spontaneous EPSP because it did not occur in other cells. The area surrounded by the red box is expanded below. **D.** EPSPs evoked by optogenetic stimuli (blue arrows) could be blocked by the AMPA receptor antagonist DNQX (10 µM; 5 GCs in 5 slices from 3 mice). A representative example of an EPSP before (Control) and after DNQX (DNQX) shows the blockade of the EPSP. **E.** EPSPs were elicited by optogenetic stimuli at 3 membrane potentials to show that at the depolarized potential an action potential was triggered. These data are consistent with weak excitation that requires depolarization of the cell to elicit firing.

### III. Paired pulse ratio

A previous study of patch-clamped GCs (Hashimotodani et al., 2017) showed weak paired-pulse facilitation. We examined whether a similar result would be obtained with our methods. For paired pulses, identical stimuli were tested with a range of interstimulus intervals (10-200 msec; Fig. 3B). Stimuli were half-maximal. The second response of the pair was usually similar or smaller than the first (Fig. 3B). Therefore, our method agreed with past studies.

### IV. Using characteristics of optogenetically-evoked fEPSPs to improve electrically-evoked fEPSPs

Paired pulse ratios were very small using 10-30 msec interstimulus intervals, consistent with the inability of ChR2 to follow high frequencies of stimulation (Wang et al., 2007). This raised the point that optogenetic methods have limitations, i.e., at high frequencies they may not be possible to use. Although faster opsins are available, there still could be future investigators who prefer electrical stimulation. Therefore we attempted to use the characteristics of optogenetic fEPSPs to modify the use of electrical stimulation of the IML so that it could more selectively activate the MC input to GCs than in the past. To this end, electrical stimulating electrodes were small in diameter (see Methods), and low currents were used to limit current spread to other layers. We placed recording electrodes in adjacent layers to ensure the responses recorded there were small relative to the fEPSPs in the IML. Moreover, we confirmed that they flipped polarity in the OML, like optogenetic fEPSPs (Fig. 1E; Supplemental Fig. 4). Last, we confirmed that fEPSPs evoked minimal population spikes in the GCL, like optogenetic fEPSPs (Fig. 3A; Supplemental Fig. 4). We found that approximately half of the slices we tested (n=8/17 slices in 10/14 mice) could meet these criteria (Supplemental Fig. 4). In those slices, paired pulse ratios showed significantly greater paired pulse ratios at 10-30 msec interstimulus intervals (Fig. 3B). In other slices, there was paired pulse facilitation, similar to characteristics of the MML input, suggesting current activated MML fibers. For those fEPSPs that met the criteria, there was little evidence of paired pulse ratios over 100% at any interstimulus interval, like optogenetically-evoked fEPSPs (Fig. 3B). Therefore, we think that use of the characteristics of optogenetically-evoked fEPSPs can improve the ability to use electrical stimulation of the IML to assess the MC input to GCs.

### V. Why optogenetic stimulation of MC axons does not evoke GC population spikes

The lack of a population spike evoked by optogenetic stimulation led us to question whether the MC input to GCs was too small to reach threshold or whether the EPSPs were inhibited by GABAergic neurons that were activated by MC axons. We first examined the monosynaptic EPSP and disynaptic IPSP produced by optogenetic stimuli. In 41 GCs, cells were depolarized to approximately −55 mV so that EPSPs would be distinguished from IPSPs. In 24/41 (54.8%) of cells there were no detectable hyperpolarizations after the EPSP. Then we returned cells to potentials between −65 and −75 mV and compared the amplitudes of the EPSP in GCs with and without IPSPs. Amplitudes were not signficantly different if there was a IPSP or not (EPSP without an IPSP, 1.89 ± 0.27 mV, n=16; EPSP with a subsequent IPSP, 1.52 ± 0.28 mV, n=10; Student’s t-test, t=0.944, df=24, p=0.354). It was surprising to find many GC EPSPs without IPSPs, but the results are consistent with the use of slices primarily from sites distant to MC somata, where MCs primarily innervate GC dendrites in the IML rather than GABAergic neurons (Buckmaster et al., 1996). Then we measured threshold in the GCs from the phase plot of an action potential evoked at threshold by a rectangular current step from RMP. Threshold was −37.54 ± 1.00 (n=43). These data support the idea that the EPSPs were not inhibited by IPSPs and were simply too small to reach threshold.

Third, we compared the MC input to the perforant path input in the same slices. We did this to determine if inhibition was simply too strong in slices to obtain population spikes. For the best comparison, we used electrical stimulation for both inputs. To do so, we took advantage of our experience to tailor the IML stimulation so it optimized activation of MC axons (as explained above). Then we adjusted stimulus intensity so the perforant path stimuli evoked comparable fEPSPs as the fEPSPs evoked by stimuli to the IML (fEPSP mean amplitude, OML vs. IML, Student’s t-test, t=0.567, df=2, p=0.601). The results showed that the perforant path stimuli evoked 2-3 mV population spikes but IML stimuli did not evoke detectable population spikes (Mann-Whitney *U* test, p=0.002).

Taken together, we suggest that the weak amplitude of the MC EPSP is the primary reason why MC-GC activation does not elicit population spikes.

### VI. Confirmation with whole-cell recording

Whole-cell recordings confirmed the results using a different recording chamber and a slightly different optogenetic method (see Methods). There were 57 whole-cell recordings of GCs in 31 mice that were 98.8 ± 2.9 days at the time of recording. Most GCs (50/57 or 87.7%) had optogenetic responses. The 50 GCs were from *Drd2*-Cre (44 cells) and the others were *Crlr*-Cre mice (6 cells). The majority of the 50 GCs were from males (34 cells) and the rest of the GCs were from females (16 cells).

Of the 50 cells with robust responses to light, all had depolarizations at RMP (Fig. 3D). In voltage clamp, EPSPs corresponded to small inward currents (Fig. 3C; n=3). Therefore, whole-cell recordings in current clamp agreed with the recordings in voltage clamp from the same cell. DNQX blocked the depolarizations (3/3 cells tested; Fig. 3D).

Quantification of EPSPs used responses recorded at a similar range of GC membrane potentials (−65 and −75 mV). Like the recordings described above, only brief optogenetic stimulation was required for responses (1.50 ± 0.11 msec; range, 0.50-2.50; n=24 cells). Also like the prior results, the latency from the onset of the light pulse to the onset of the EPSP was consistent with a monosynaptic pathway (3.13 ± 0.12 msec, range, 2.40-4.10; n=16 cells). Measured from the end of the light pulse to the onset of the EPSP, the latency to onset of the EPSP was 1.62 ± 0.17 (range, 0.40-3.04; n=16 cells). The mean peak amplitude of EPSPs was small (1.89 ± 0.27 mV; range, 0.39 - 3.60 mV; n=16 cells) but depolarization of GCs could elicit APs (Fig. 3E).

### VII. Effects of the septotemporal axis

#### A. Slice location

The hippocampus was divided into a dorsal, mid and ventral region, defining each part as one-third of the hippocampus (Fig. 4A). Data from all mice were pooled. For fEPSPs, a two-way ANOVA was conducted with slice location and type of measurement (slope or amplitude) as main factors. There was no effect of slice location (Fig. 4B). EPSPs in whole-cell recordings were also analyzed and they confirmed there was no effect of slice location (Fig 4B). Thus, fEPSPs were not significantly different in dorsal, mid or ventral regions.

**Figure 4.**
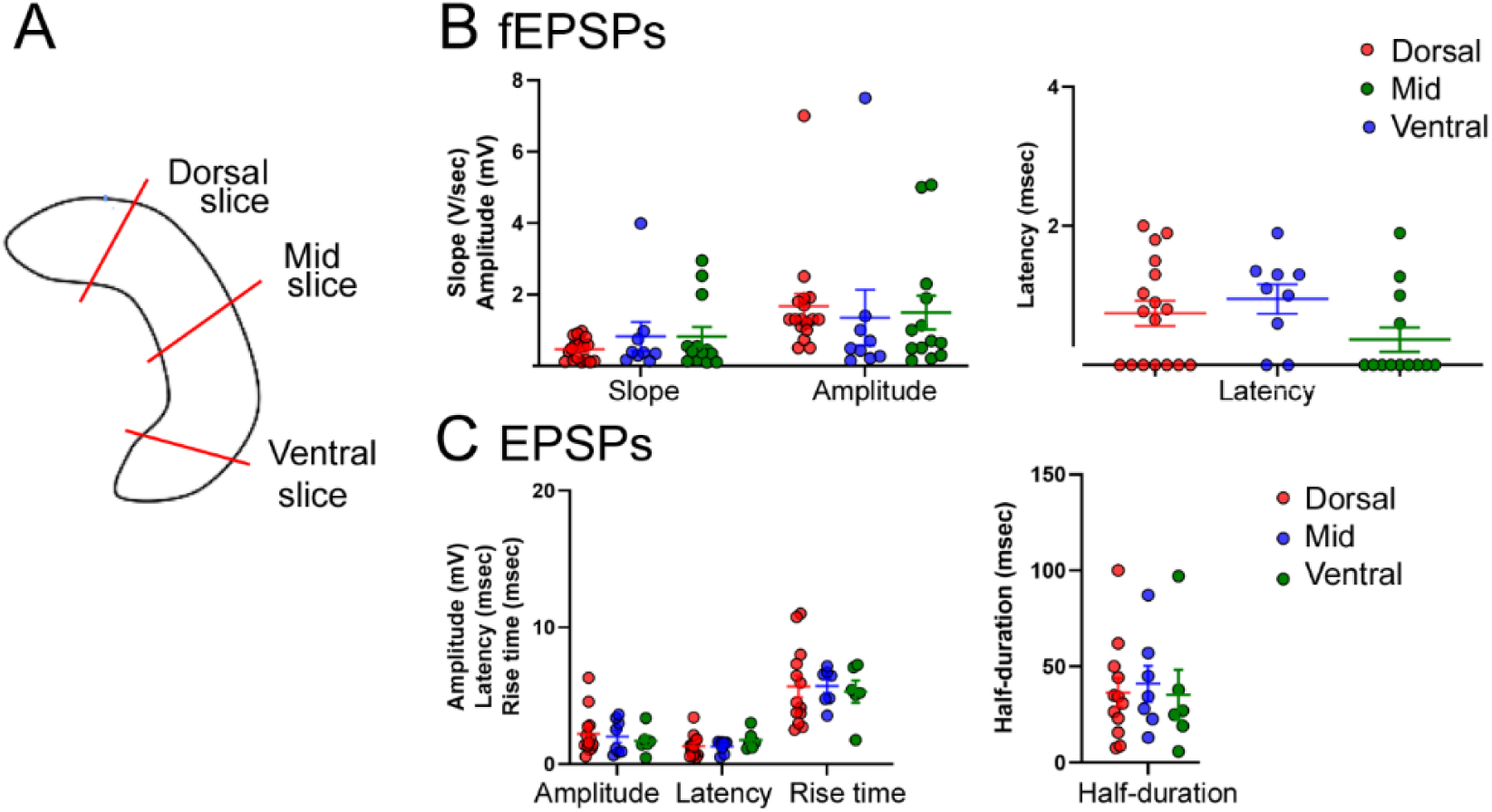
Similarity of optogenetically-evoked fEPSPs across the septotemporal axis. A. A diagram showed where slices were selected for the comparisons in B and C. B. A two-way ANOVA was conducted with slice location and type of measurement (slope or amplitude) as main factors. There was no effect of slice location (F(2,72)=0.03; p=0.969). C. EPSPs in whole-cell recordings confirmed that there was no effect of slice location (two-way ANOVA, F(2,93)=0.10; p=0.905).

### B. Dorsal vs. ventral viral injection sites

Slices were compared for animals injected in dorsal or ventral DG (Fig. 5A1). All slices from these animals were included, whether they were near or far from the injection site. There was no effect of the injection site location on fEPSP slope or amplitude (Fig. 5A2). Latencies of fEPSPs also were not different (Fig. 5A2). These data suggest similarity across the septotemporal axis.

**Figure 5.**
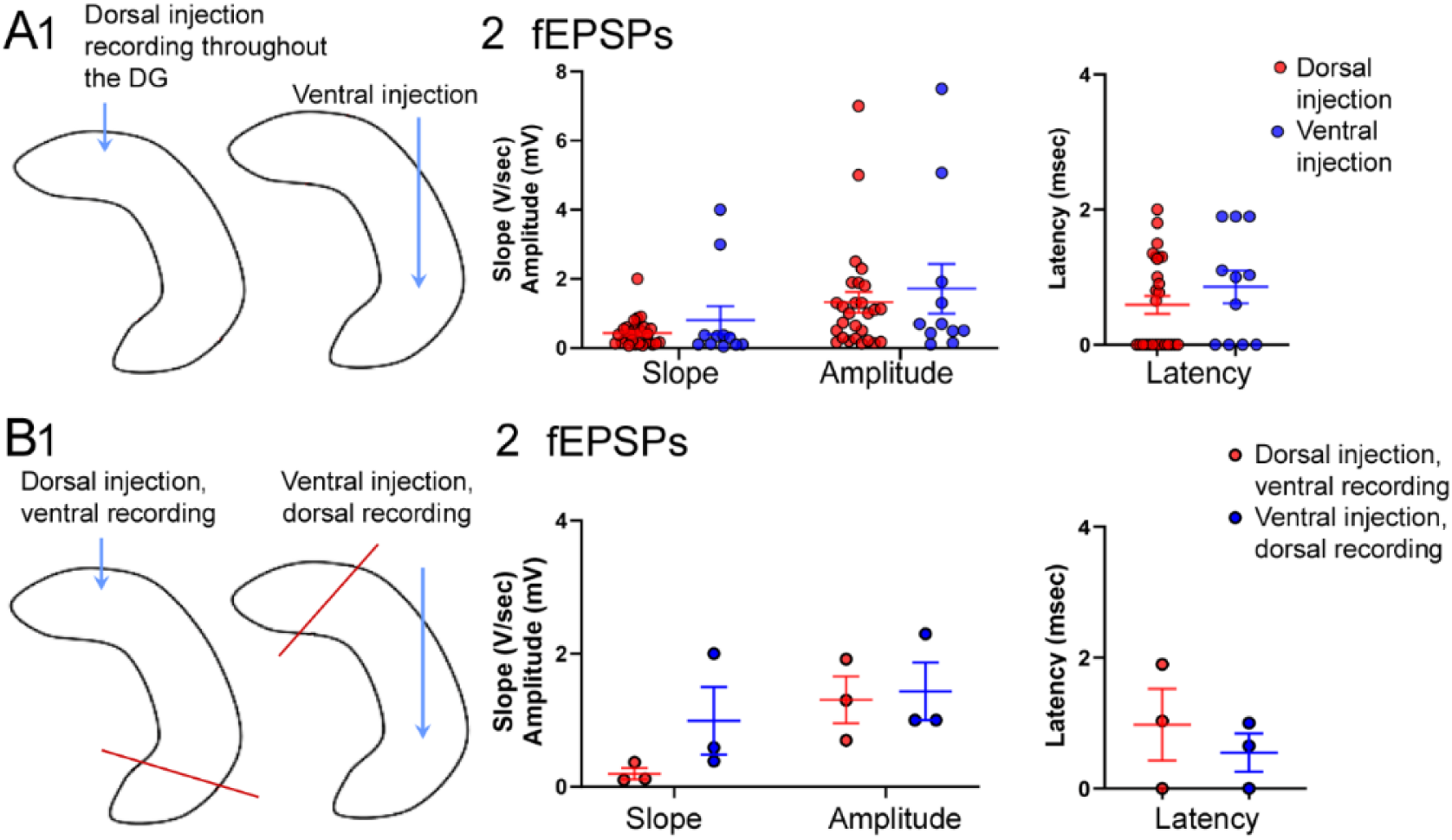
Optogenetic responses of GCs were similar for dorsal and ventral MC axons. A. 1. A diagram shows a dorsal injection site compared to a ventral injection. Only one injection was made, either dorsal or ventral. Slices were pooled from all regions of the DG. 2. A two-way ANOVA showed no effect of injection site location (F(1,72)=0.0005; p=0.981). Latencies of fEPSPs also were not different (Mann-Whitney *U* test, p=0.350; *U*, 120.5). B. 1. In different animals, an Injection was either made in the dorsal DG and slices were selected from the ipsilateral ventral DG or a ventral injection was made and slices were selected from the ipsilateral dorsal DG. 2. A two-way ANOVA showed no effect of dorsal injections vs. ventral injections on slope or amplitude (F(1,4)=-0.88, p=0.401). There were no differences in latency (Mann-Whitney *U* test, p=0.500, *U*, 2.50).

As mentioned above, the distant ipsilateral axons of dorsal MCs extend not only in the IML but also other subdivisions of the ML. However, the distal ipsilateral axons of ventral MCs are restricted to the dorsal IML. Therefore, we either injected dorsally and selected slices in the ventral DG or injected ventrally and made slices from dorsal DG (Fig. 5B1), hypothesizing that fEPSPs would differ. However, the results showed no detectable differences in optogenetic responses (Fig. 5B2). Note that these results do not necessarily conflict with prior results suggesting ventral MCs synergize more with the PP input or that ventral MCs might be more excitatory than dorsal MCs but suggest that there is little difference in fEPSPs produced by dorsal and ventral MCs.

#### C. Ipsilateral vs. contralateral recording sites

Recording sites that were ipsilateral to the injection site were divided into those that were close to it (“near”) or far from the injection site (“far”). The definitions for ‘near’ or ‘far’ slices were based on studying ChR2 expression and finding that slices at least 800 µm away from the injection site had few (<5) MC somata expressing ChR2. Therefore these slices were considered far from the injection site. In contrast, slices at the injection site or adjacent to it were called “near” (Fig. 6A).

**Figure 6.**
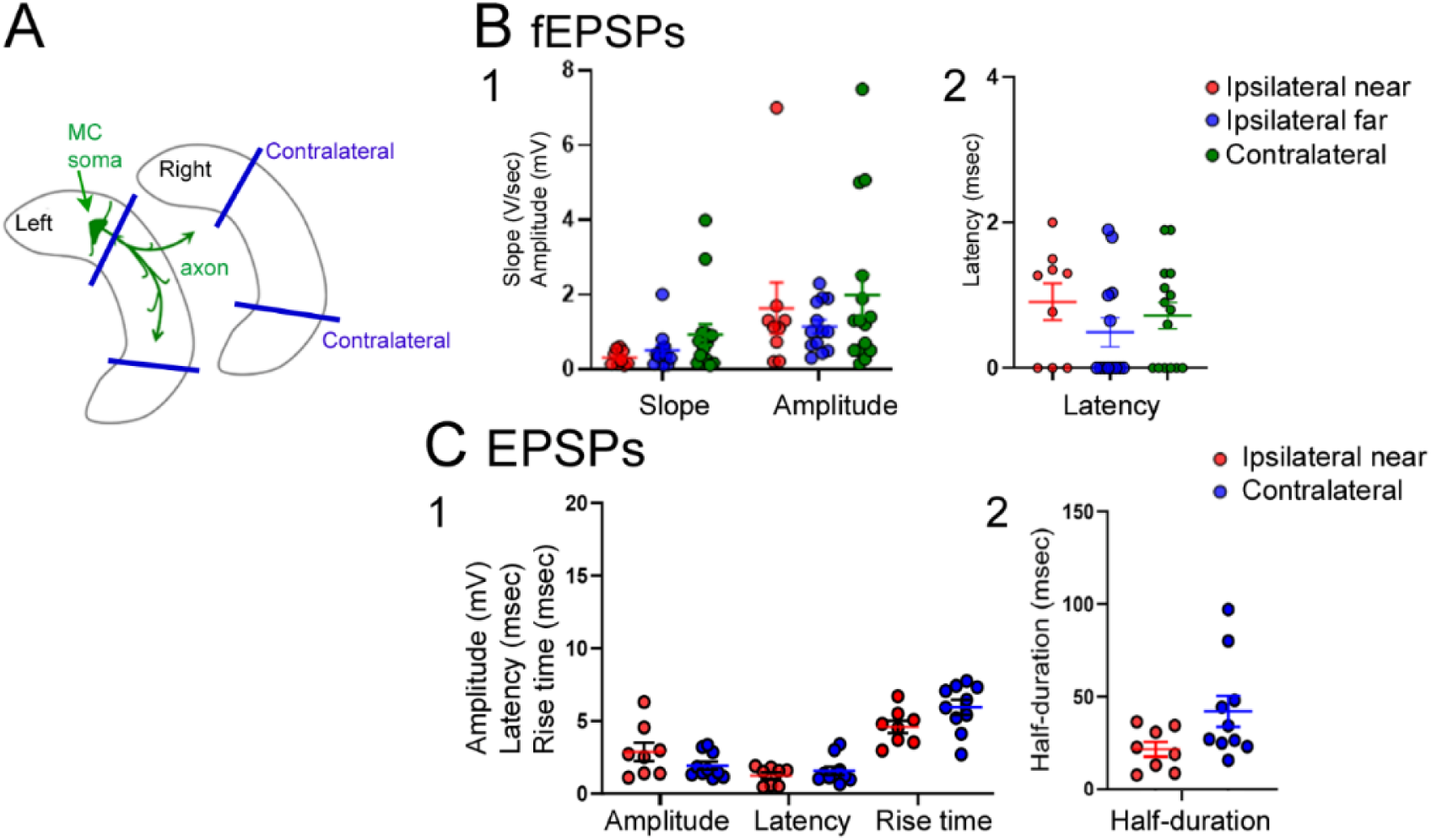
Optogenetic responses were similar between slices ipsilateral and contralateral to viral injection. A. A diagram shows how slices were selected for a dorsal injection into the left DG. Slices were either near the injection (“ipsilateral near”), far but still in the ipsilateral hippocampus (“ipsilateral far”) or contralateral. “Far” was defined as > 800 µm from the injection site because at this distance from the injection site few MCs expressed virus B 1. There were no significant differences in fEPSP slope (Kruskal-Wallis ANOVA; p=0.126; Kruskal-Wallis statistic, 4.14) or amplitude (p=0.302; Kruskal-Wallis statistic 2.40). 2. The same was true for latency to onset (p=0.830; Kruskal-Wallis statistic 0.37). C. Whole-cell recordings were used as a confirmation, but only ipsilateral “near” and contralateral were compared because of a small sample size for slices that were ipsilateral “far”. A two-way ANOVA showed no effect of location on EPSP amplitude, latency or rise time (1) or half-duration (F(1,17)=0.909; p=0.354).

Ipsilateral “near”, ipsilateral “far”, and contralateral recording sites were compared (Fig. 6A). There were no significant differences in fEPSP slope, amplitude or latency to onset (p=0.830; Kruskal-Wallis statistic 0.37; Fig. 6B1-2).

Whole-cell recordings were used to ask if the results would be the same, but only ipsilateral “near” and contralateral were compared because of a small sample size for slices that were ipsilateral “far”. There was no effect of location (Fig. 6C).

#### D. Recording location in the dentate gyrus blades

The GCL was divided into three parts, the superior blade, the crest and the inferior blade (Supplemental Fig. 5A). Statistical comparisons were restricted to the superior blade and crest because optogenetic stimuli were tested in the lower blade in very few slices. There was no significant difference between the GCs in the superior blade and crest for slope, amplitude, or latency (Supplemental Fig. 5B).

For EPSPs recorded in whole-cell mode, a two-way-ANOVA was conducted with location (superior blade or crest) and type of measurement (EPSP amplitude, latency to onset, rise time and half-duration) as main factors. There was no effect of location (Supplemental Fig. 4C). The same percentage of EPSPs were recorded in the superior blade (11/21 or 52%) and crest (10/21 or 48%; Fisher’s exact test, p>0.999).

#### E. Effect of sex

There were no detectable sex differences in fEPSPs or EPSPs (Supplemental Fig. 6). When fEPSPs from Drd2-Cre and Crlr-Cre mice were compared, there were no significant differences in fEPSP slope (Student’s t-test, t=1.13, df 33, p=0.265) or amplitude (Mann-Whitney *U* test, p=0.702; *U* statistic 135) in the two mouse lines (data not shown). Latencies to onset of fEPSPs were not significantly different (Mann-Whitney *U* test, p=0.929; *U* statistic 158; data not shown). There were no detectable differences in the number of slices with and without detectable optogenetic responses (Fisher’s exact test, p=0.337; data not shown).

## DISCUSSION

### I. Summary

The results provide methods to use extracellular recordings in hippocampal slices to ask questions about the MC input to GCs with specificity. This method enables opportunities to ask questions about this pathway without requiring more advanced training or materials. Although there are advantages to more sophisticated techniques, there are benefits to an easier approach, such as ease of use for a non-specialist. There also are benefits to extracellular recordings, such as sampling of subsets of neurons rather that one neuron at a time. Moreover, patch clamping involves dialysis of the recorded cell, which has drawbacks.

This study showed that MCs have weak excitatory effects on GCs, which has been suggested before based on other methods. We found excitatory effects on GCs were common across all parts of the septotemporal axis, which has not been shown before. The widespread excitation of GCs, even if weak, would make MCs able to exert a widespread influence, especially if they were activated synchronously.

Although the relatively small fEPSPs that we recorded might be doubted without confirmation by other methods, the agreement with data from whole-cell recordings provided support for the findings.

### II. Relevance to past studies of MCs

The common nature of excitation was predicted by anatomical observations (Buckmaster et al., 1992; Buckmaster and Schwartzkroin, 1994; Buckmaster et al., 1996). However, anatomical studies have shown that MCs contact GABAergic neurons also, and several studies have suggested that MCs mainly inhibit GCs for this reason. Therefore the common nature of excitation was somewhat surprising. On the other hand, many patch clamp recordings of GCs show that MC excitation is present without subsequent IPSCs (Chancey et al., 2014) or before inhibition of GCs (Hashimotodani et al., 2017). Moreover, these recordings sometimes show excitatory currents that are as large or larger than inhibitory currents (Hashimotodani et al., 2017).

### III. EPSP homogeneity

A remarkable finding was the consistency of fEPSPs across the septotemporal axis. This is surprising because there are differences in dorsal and ventral MC expression of numerous peptides (Cembrowski wt al., 2016). In addition, dorsal and ventral MCs show electrophysiological differences (Jinno et al., 2003). Ventral MCs also express more c-fos than dorsal MCs in response to novelty (Duffy et al., 2013; Bernstein et al., 2019) and influence novelty detection (Fredes et al., 2020). These results may be true, but the dorsal and ventral terminals of MCs seem to have similar effects when examined in slices. This idea is consistent with early theories of hippocampus as a structure made of repeating units or lamella (Andersen et al., 2000).

### IV. General pros and cons of the method

The approach described here, using optogenetics and viral delivery of opsins to assay the MC input to GCs has several advantages and disadvantages.

An advantage is the precision of optogenetics with viral delivery of Cre-dependent virus expressing opsins. However, control experiments were necessary to be sure there non-specific effects of light were absent, and the expression was specific. Also, the volume of virus to inject requires pilot experiments because high volumes might lead to a loss of specificity but low volumes may not lead to robust expression. Furthermore, the time after injection when expression is robust needs to be checked and if long periods of time are required this can delay experiments. We found that delays of 5-7 days led only to expression in somata and dendrites of MCs, but delays 14-30 days led to robust expression of axons ipsilateral and contralateral to the injection site.

There are some electrophysiological limitations of optogenetics compared to electrical stimulation. For optogenetic activation of an input, one can not define the precise onset of responses since it is not clear when, during the msec of light stimulation, presynaptic fibers become active. For short delays to a response, such as monosynaptic delays, this problem may be significant. We circumvented the problem by measuring time to onset of a fEPSP or EPSP in two ways, one from the start and one from the end of the light pulse, but the limitation is not entirely mitigated by that approach. Optogenetics also is difficult to use with high frequency trains because opsins like ChR2 do not follow high frequencies. On the other hand, fast opsins can avoid this limitation. Alternatively, high frequencies can be tested with electrical stimuli, given we showed a way to use what we learned with optogenetics to tailor the use of electrical stimulation to activate the MC input to GCs. Electrical stimulation also can be useful because it is less costly and time-consuming than optogenetics and requires no viral injection.

An advantage of the method we describe is the utility to assay the MC-GC pathway with extracellular recordings, because this technique requires less training than whole-cell recordings. Another advantage is the ability to use the approach with interface as well as submerged chambers, since the interface chamber has advantages over the submerged chamber and vice-versa. One advantage of the interface chamber is that many slices from one mouse can be sampled together, allowing within-mouse comparisons. Although submerged slices can be compared one at a time, the slices that are evaluated later in the day were placed in a holding chamber for a long time, and that may lead to different results. On the other hand, the submerged chamber allows visualization of the DG layers, providing the opportunity to determine the viability of the cells. With an interface chamber, a stimulus must be used to screen slices to determine they are healthy prior to use.

Using an interface chamber, the optic fiber can be placed in the air above the slice where it does not damage the slice; for submerged chambers our optic fiber penetrated the fluid over the slice. We have not tested our method with the light originating from an objective in a submerged slice, but others do so routinely.

### V. Pros and cons of using slices

A disadvantage of an approach that uses slices is that behavioral states such as sleep are not possible to examine. On the other hand, numerous investigators have simulated sleep in slices by adding pharmacological agents that naturally rise during sleep, such as the cholinergic agonist carbachol (Colombi et al., 2016), or added neurotransmitters that promote arousal to simulate wakefulness (Hinard et al., 2012).

An advantage of in vivo recordings is the ability to keep the entire MC intact. However, this advantage is debatable because cut projections have been studied for decades in slices and a lot of the data are similar to those obtained in vivo. Moreover, in order to take advantage of the intact MC projection in vivo one needs to define the subdivisions of the ML which is not easy when they can not be visualized. In addtion, in vivo studies of temporal DG are more difficult than dorsal DG because of the depth of ventral DG. Perhaps for this reason many studies of MCs in vivo are based only on recordings from dorsal DG (Sloviter, 1991a, b; Hsu et al., 2015).

A disadvantage of some in vivo studies is the reliance on unit recordings from GCs. Although this approach had led to a wealth of useful data, for inputs that appear to be mainly subthreshold, like the MC input to GCs, unit firing alone will lead to less information about the MC input to GCs. One could argue that this input can not be important if it is usually subthreshold. However, a counterargument is that under some conditions the MC input to GCs may become able to cause APs in GCs. These conditions include synchronization of MCs by sharp-wave ripples, states of reduced GABAergic inhibition like sleep, or conditions that reduce endocannabinoid levels which would decrease the normal suppression of MC glutamate release on GCs (Chiu and Castillo, 2008).

### VI. Conclusions

Here we show a method to assay MC input to GCs by extracellular recordings in slices. We confirm that this method successfully recapitulates several known characteristics of the input, such as its weak but common effects on GCs, weak paired pulse ratios, and the ability of optogenetic MC activation to trigger disynaptic inhibition of GCs. We also show how to use the method to adapt electrical stimulation to make it more specific. Additionally, we report a remarkable homogeneity of the MC input across the septotemporal axis and sex which to our knowledge has not yet been addressed.

## Acknowledgements

We thank Karl Deisseroth and Charu Ramakrishnan for their help in this study. This project was supported by NIH F30 MH-110165 (HLB), NIH R01 MH-109305 NIH R01 MH-109305, NIH R37 NS-126529 (HES), the Savoy Foundation (JJB) and NSERC (JJB), and the New York State Office of Mental Health (HES).

## FIGURE LEGENDS

**Supplemental Table 1.**
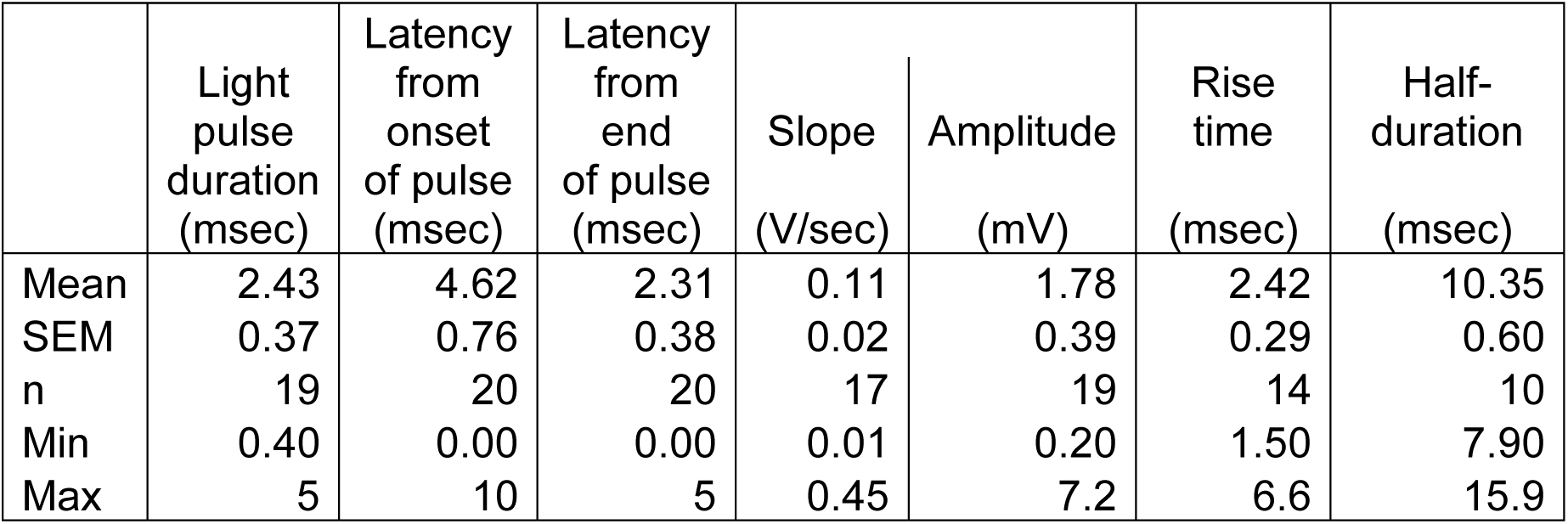
Characteristics of fEPSPs. Measurements of fEPSPs are listed using a stimulus that elicited a fEPSP that was close to the peak amplitude. The light pulse duration was the duration of the optogenetic stimulus used to elicit fEPSPs. Latency to onset was the time between the light pulse and the onset of the fEPSP, measured either from the start of the light pulse or the end. Peak amplitude was measured from the baseline just before the stimulus and was measured to the fEPSP peak. The rise time was the duration of the initial (rising) phase and was measured from the fEPSP onset to peak. The half-duration was the time between the end of the light pulse to the point on the decay of the fEPSP when it reached half its peak amplitude.

|  | Light pulse duration (msec) | Latency from onset of pulse (msec) | Latency from end of pulse (msec) | Slope (V/sec) | Amplitude (mV) | Rise time (msec) | Half-duration (msec) |
| --- | --- | --- | --- | --- | --- | --- | --- |
| Mean | 2.43 | 4.62 | 2.31 | 0.11 | 1.78 | 2.42 | 10.35 |
| SEM | 0.37 | 0.76 | 0.38 | 0.02 | 0.39 | 0.29 | 0.60 |
| n | 19 | 20 | 20 | 17 | 19 | 14 | 10 |
| Min | 0.40 | 0.00 | 0.00 | 0.01 | 0.20 | 1.50 | 7.90 |
| Max | 5 | 10 | 5 | 0.45 | 7.2 | 6.6 | 15.9 |

**Supplemental Figure 1.**
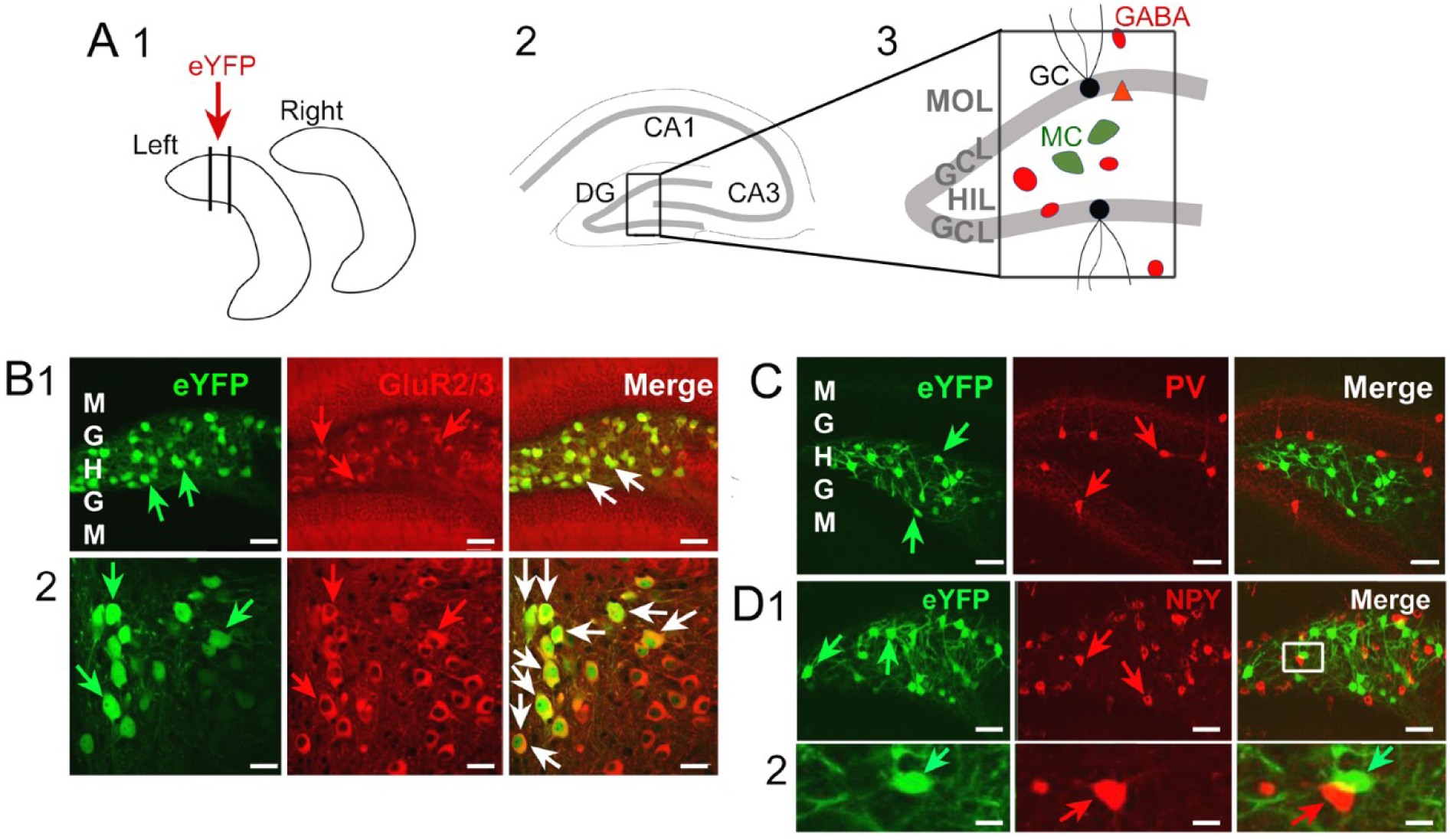
Specificity of Cre expression in MCs. **A** 1. A schematic illustrates the injection of AAV2-eYFP in the left dorsal hilus. Sections were taken for B-D from the area near the injection site 2-3. The orientation of the tissue sections in B-D. The area outlined by the box in 3) is shown in B-D. **B** 1. Tissue sections were processed using an antibody to GluR2/3 to label glutamatergic neurons. *Left:* EYFP-expressing cells are denoted by the green arrows. *Center:* GluR2/3 labels glutamatergic cells in the hilus (red arrows), most of which are MCs. *Right:* The merged image of eYFP and GluR2/3 immunofluorescence shows that eYFP-expressing hilar cells also expressed GluR2/3 (white arrows). These data suggest that the hilar cells expressing Cre were MCs. Calibration, 50 µm. M, molecular layer; G, granule cell layer; H, hilus. 2. The same sections as B1 are shown at higher gain. Calibration, 20 µm. **C.** Different sections were stained with an antibody to parvalbumin (PV), a marker of a major class of DG GABAergic neurons *Left:* eYFP-fluorescent hilar cells (green arrows) *Center:* PV-immunoreactive cells (red arrows) *Right:* The merged image shows no double-labeling. Calibration, 50 μm. **D** 1. An antibody to neuropeptide Y (NPY) was used to stain a major subset of GABAergic hilar cells. *Left and Center*: Many hilar neurons expressed eYFP (green arrows) or NPY (red arrows). *Right:* Yellow cells were rare. One area of the hilus with yellow is expanded in D2. Calibration, 20 µm. 2. The area outlined by the box in D1 is shown at higher gain to illustrate that yellow reflected overlapping cells but not double-labeling *Left:* A cell expressing eYFP is noted by the green arrow. *Center:* The red arrow points to a cell with NPY immunofluorescence that is nearby. *Right:* The merged image shows that the cells denoted by the arrows overlap and therefore yellow is observed, but they are two different cells. Calibration, 10 µm.

**Supplemental Figure 2.**
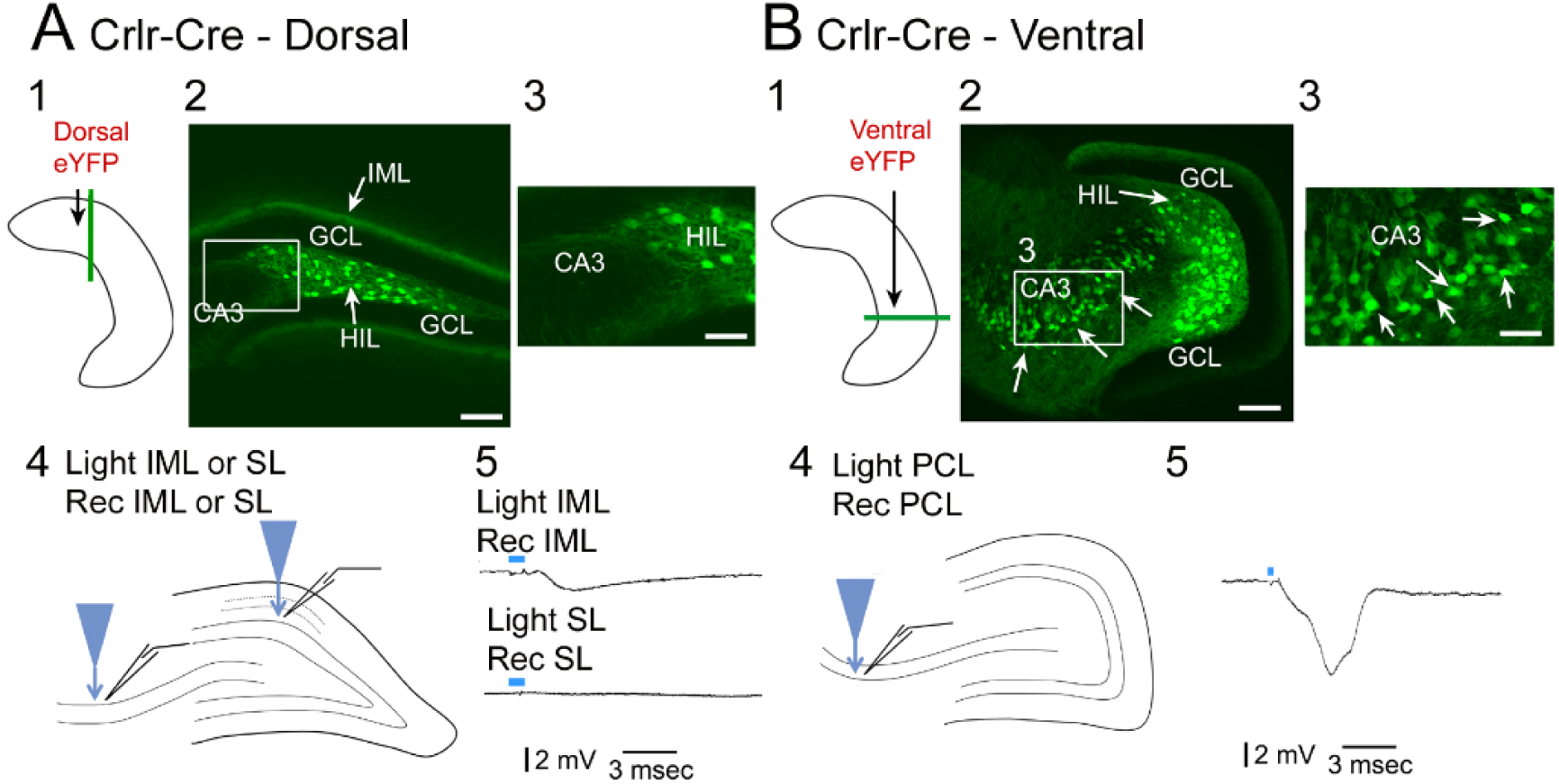
Limitations of Crlr-Cre mice. **A.** 1. A schematic shows dorsal injection of AAV2-eYFP to evaluate specificity for MCs in dorsal hippocampus. 2. After dorsal injection of AAV2-eYFP, coronal sections were made of the dorsal HC. A dorsal coronal section shows hilar fluorescence as well as IML fluorescence (arrows), indicating labeling of MCs. The area surrounded by the box is the border of CA3 and the hilus and is shown at higher gain in 3. 3. Area CA3c showed no fluorescent cells. Calibrations:100 µm (2), 50 µm (3). 4. A schematic illustrates recording sites in slices that were prepared approximately two weeks after AAV2-ChR2 injection. The optical fiber was either directed towards the IML and recordings were made there, or light was directed at the stratum lucidum (SL) in CA3 and recordings were made there. 5. *Top:* A light pulse (blue bar) directed to the IML elicited a typical fEPSP in the IML. *Bottom:* A light pulse to SL did not elicit a response. Similar data were elicited from 4 other slices (4 additional mice, i.e., 1 slice/mouse). **B.** 1. A schematic illustrates the approach to inject AAV2-eYFP into ventral hippocampus. After ventral injection, horizontal sections were made of the ventral HC. 2-3. A ventral horizontal section shows hilar fluorescence as well as weak IML fluorescence. The area surrounded by the box is shown at higher gain in 3. Arrows point to fluorescent CA3 neurons. Calibrations, 100 μm (2); 50 μm (3). 4. A schematic illustrates the light fiber direction towards the CA3 cell layer and recording at that location. 5. A robust light response was evoked with a short latency. These data suggest a lack of specificity for MCs in ventral hippocampus of Crlr-Cre mice.

**Supplemental Fig. 3.**
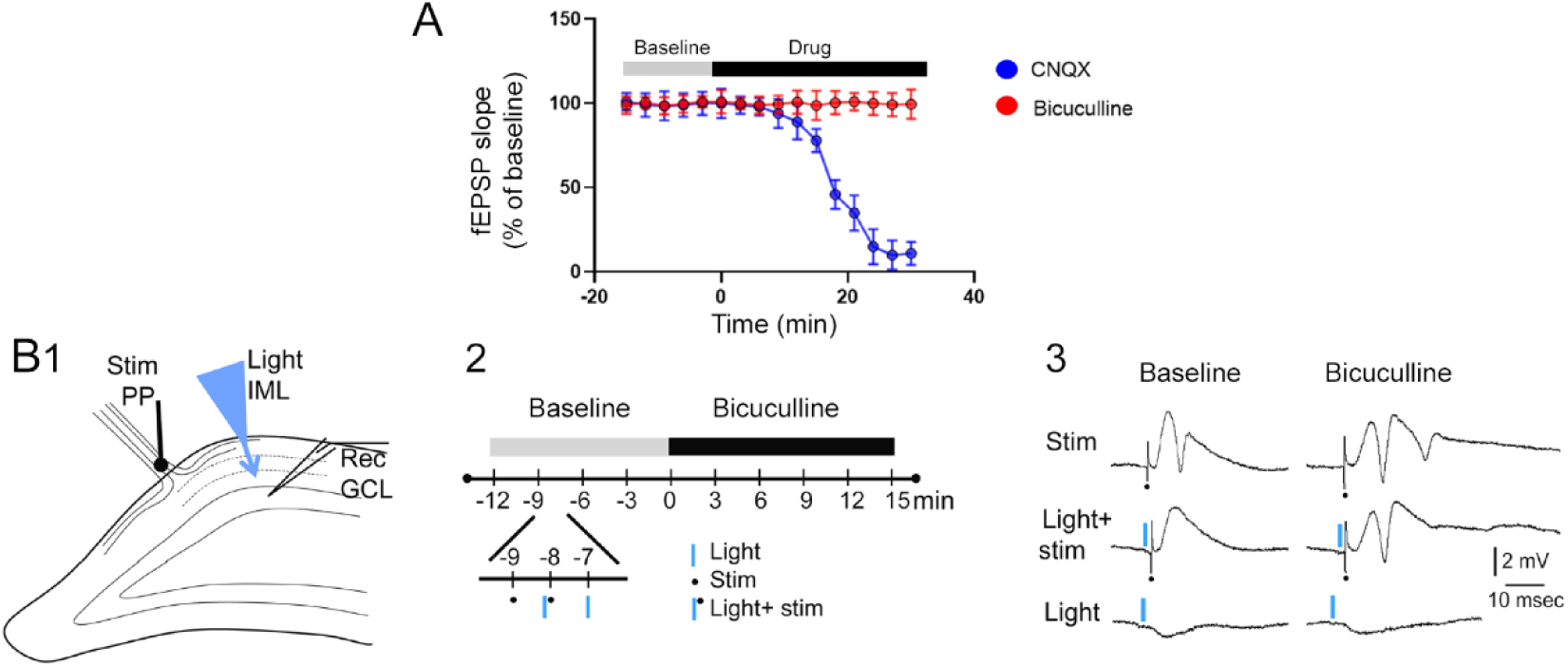
AMPA receptors, but not GABA_A_ receptors, contribute to MC→GC optogenetic excitation. **A.** Optogenetic stimuli were triggered to evoke fEPSPs. FEPSPs decreased in amplitude upon exposure to 10 µM CNQX (3 slices, 3 mice) but not 10 µM BMI (3 different slices, 3 different mice). **B** 1. A schematic of the recording arrangement. 2. A schematic of the experimental timeline. Stimuli were triggered every minute in this order: electrical stimulation of the PP, optogenetic stimulation, both PP and optogenetic stimulation. For the latter, the optogenetic stimulus was triggered with the same delay (the pulse was 1 msec long, and started 2 msec before the PP stimulus). This sequence was repeated throughout the experiment. After a baseline period to determine that responses were stable, the slice was exposed to 10 µM BMI. 3. Responses are shown during the baseline and after BMI was added to the aCSF. In the baseline period, a PP stimulus (Baseline, Stim) evoked a large population spike and a prepulse of light inhibited it (Baseline, Light + Stim). When light was triggered by itself (Baseline, Light), a small fEPSP occurred. After BMI, the light pulse no longer inhibited the population spike elicited by PP stimulation (BMI, Light+ Stim). The fEPSP evoked by the light pulse appeared to be unaffected by BMI (BMI, Light), which is consistent with the idea that optogenetic stimulation evoked a fEPSP mediated by glutamate, not GABA. Note that the second population spike evoked in the presence of BMI (BMI, Stim), was inhibited by the light pulse (BMI, Light + Stim). This result is consistent with BMI-resistant GABA_B_ receptors causing a slow inhibition of the response to the PP. Results were replicated in 5 slices from 5 mice.

**Supplemental Fig. 4.**
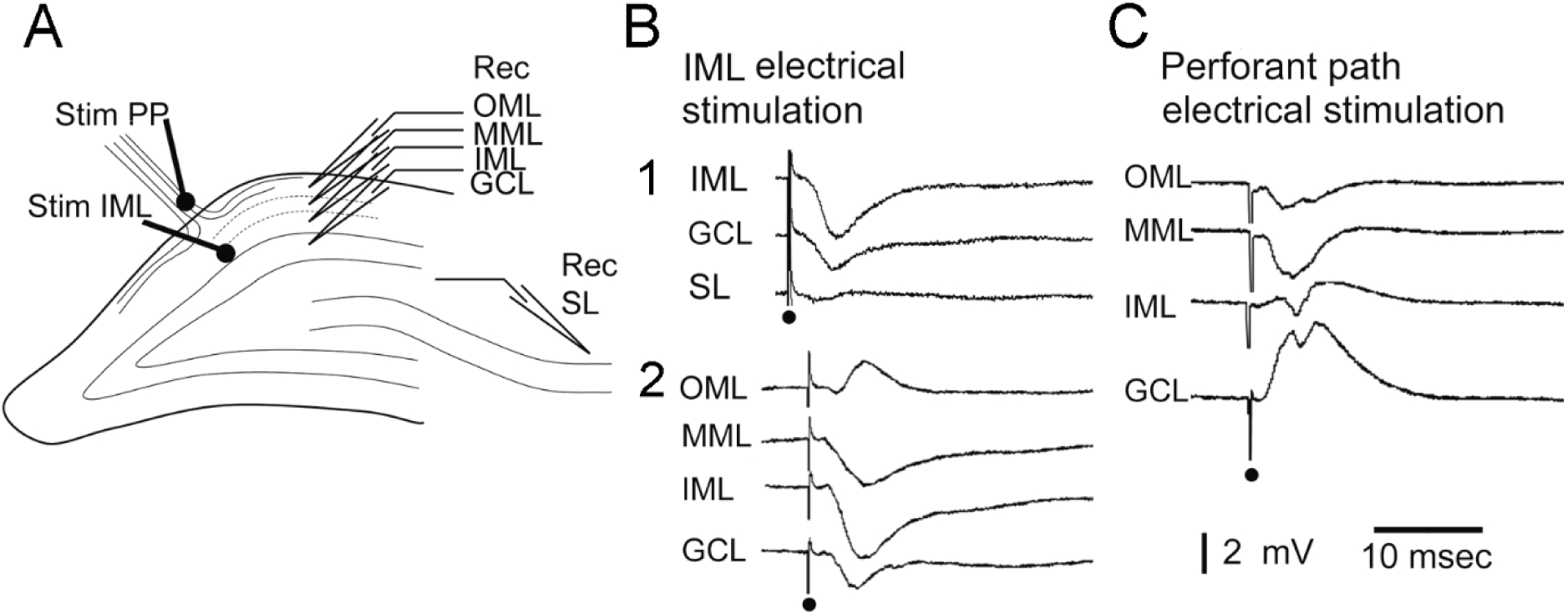
Electrical stimulation of the IML can be used to evoke fEPSPs like those produced by optogenetic activation of MC axons. **A.** A diagram shows the recording site in the GCL and the stimulating electrode in the PP. The site of stimulation was on the hippocampal fissure rather than the ML to avoid direct stimulation of DG neurons. Light was directed to the IML near the GCL recording site. Electrical stimulation is not selective for MC axons, so to make electrical stimulation as specific as possible for the IML we used a stimulus electrode that was small in diameter (25 µm diameter) and weak currents (10-50 µsec, 50 µA). In this way, current would not be likely to spread to the MML and activate PP axons. In addition, we only used slices where an electrical stimulus to the IML evoked a fEPSP that was larger in the IML than the MML. **B.** 1. Example of a fEPSP elicited by electrical stimulation of the IML. Sequential recordings in different layers showed the largest responses in the IML, consistent with preferential stimulation of the IML axons. Note there is no population spike, and no response in the SL of CA3, consistent with weak activation of the IML input to GCs. 2. Another example of fEPSPs elicited by IML electrical stimulation, demonstrating IML axons were preferentially stimulated, because the fEPSP that was maximal in the IML was small in the MML and flipped polarity in the OML. **C.** In contrast to IML stimulation, electrical stimulation of the PP elicited a fEPSP that could trigger a population spike in the GCL. Same slice as B. The PP stimulus strength was lowered so the fEPSP was comparable in amplitude to the fEPSP generated in the IML by IML stimulation. These data suggest that the MC input to GCs is weakly excitatory compared to the PP input.

**Supplemental Figure 5.**
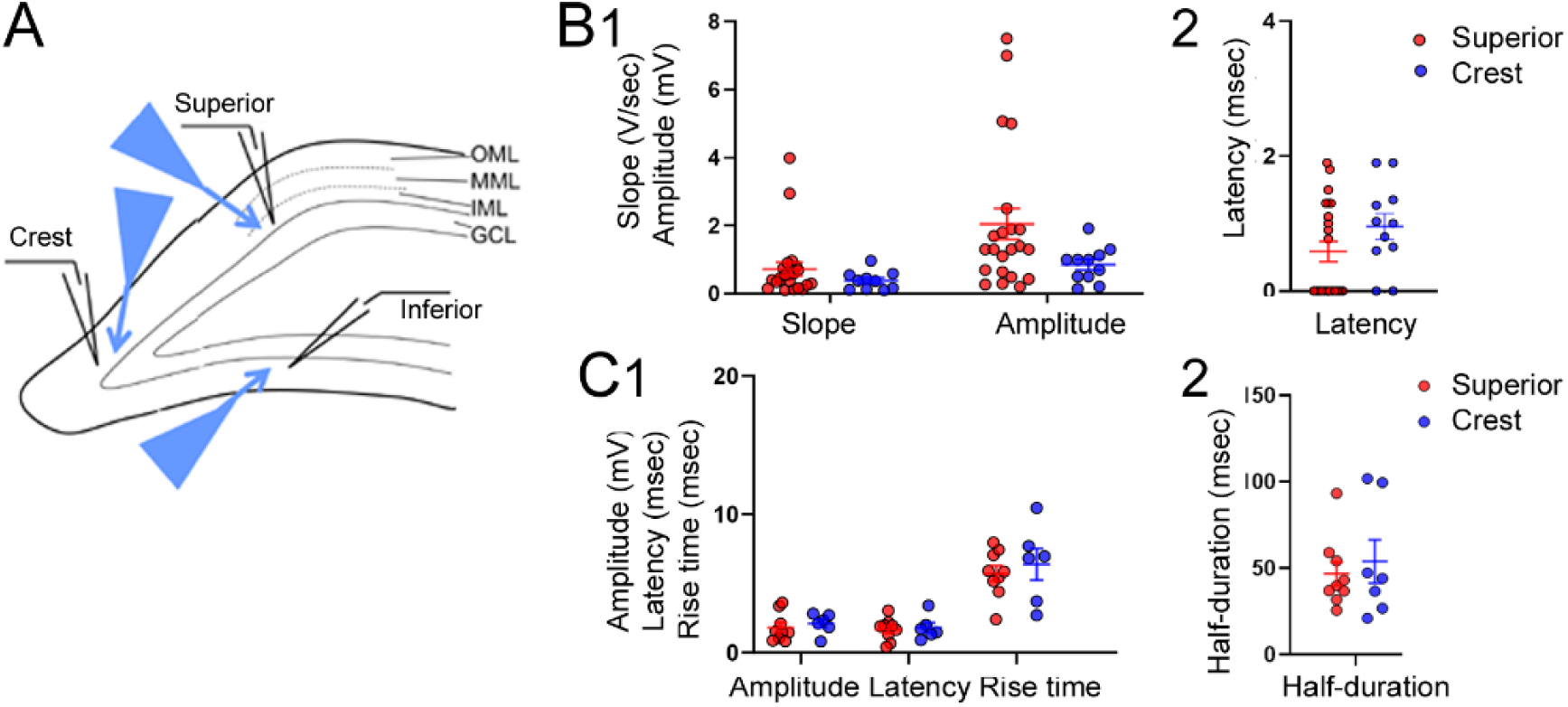
Comparisons were made of optogenetic responses in the different parts of the DG blades using either coronal or horizontal orientations. **A.** The upper blade, lower blade and crest regions are demarcated as in prior studies (Scharfman, 1995). There were few samples of the lower blade so statistical comparisons were made for the superior blade and crest in B and C. **B.** 1. There were no significant differences between the GCs in the superior blade and crest for slope (Mann-Whitney *U* test, p=0.157, *U* =83.5) or amplitude (p=0.070; *U*=73.5) 2. There were no significant differences in latency (p=0.164; *U*=85.5). **C.** For EPSPs, there was no effect of location on amplitude, latency to onset, rise time (1) or half-duration (2; two-way ANOVA, (F(1,52)=0.0004, p=0.984).

**Supplemental Fig. 6.**
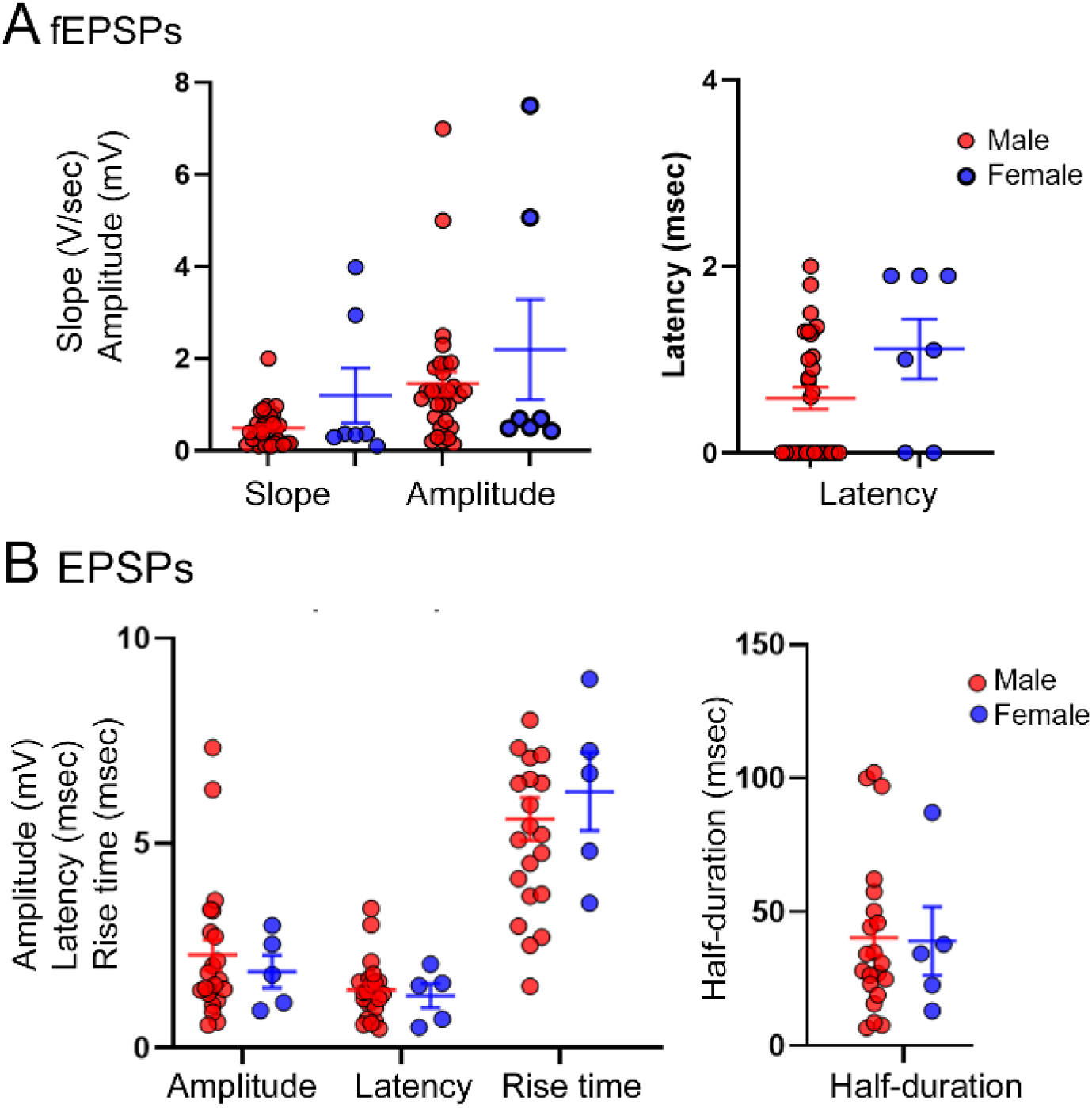
No effects of sex on fEPSPs evoked by optogenetic stimuli. **A.** FEPSPs were studied in 11 male mice (30 slices) and 3 female mice (7 slices). A two-way ANOVA showed no main effect of sex (F(1,70)=0.29; p=0.589). **B.** EPSPs were compared using whole-cell recordings at −65 to −75 mV. A two-way ANOVA was conducted with sex and EPSP measurement (amplitude, latency to onset, rise time, or half-duration) as main factors. There was no significant effect of sex (F(3,99)=0.24; p=0.866).

